# Dynamic Expression of Brain Functional Systems Disclosed by Fine-Scale Analysis of Edge Time Series

**DOI:** 10.1101/2020.08.23.263541

**Authors:** Olaf Sporns, Joshua Faskowitz, Andreia Sofia Teixera, Richard F. Betzel

**Affiliations:** Department of Psychological and Brain Sciences, Indiana University, Bloomington, USA; Program in Neuroscience, Indiana University, Bloomington, USA; Network Science Institute, Indiana University, Bloomington, USA; Cognitive Science Program, Indiana University, Bloomington, USA; Center for Social and Biomedical Complexity, School of Informatics, Computing, & Engineering, Indiana University, Bloomington IN, USA; INESC-ID, Lisboa, Portugal

## Abstract

Functional connectivity (FC) describes the statistical dependence between brain regions in resting-state fMRI studies and is usually estimated as the Pearson correlation of time courses. Clustering reveals densely coupled sets of regions constituting a set of resting-state networks or functional systems. These systems manifest most clearly when FC is sampled over longer epochs lasting many minutes but appear to fluctuate on shorter time scales. Here, we propose a new approach to track these temporal fluctuations. Un-wrapping FC signal correlations yields pairwise co-fluctuation time series, one for each node pair/edge, and reveals fine-scale dynamics across the network. Co-fluctuations partition the network, at each time step, into exactly two communities. Sampled over time, the overlay of these bipartitions, a binary decomposition of the original time series, very closely approximates functional connectivity. Bipartitions exhibit characteristic spatiotemporal patterns that are reproducible across participants and imaging sessions and disclose fine-scale profiles of the time-varying levels of expression of functional systems. Our findings document that functional systems appear transiently and intermittently, and that FC results from the overlay of many variable instances of system expression. Potential applications of this decomposition of functional connectivity into a set of binary patterns are discussed.

## Introduction

Modern network neuroscience conceptualizes the brain as an interconnected dynamic multiscale system (Bullmore and Sporns, 2009; Bassett and Sporns 2017; Betzel and Bassett, 2017). At the level of the whole brain, anatomical projections between brain regions shape spontaneous dynamics and constrain the brain’s momentary responses to changes in input, internal state, and environmental demand (Honey et al. 2009; Suárez et al. 2020). Statistical dependencies among regional time courses are described as ‘functional connectivity’, quantified with a variety of bivariate metrics (Friston 2011; Buckner et al 2013). In extant fMRI research, the Pearson correlation of blood oxygenation level dependent (BOLD) time courses remains in wide use, generally applied to long epochs of resting or task-evoked responses. The resulting correlation matrix, representing a functional network (Power et al 2011) or ‘functional connectome’ (Biswal et al. 2010), provides a summary representation of the system’s pairwise dependencies.

Functional connectivity, measured during the resting-state, exhibits highly consistent patterns across imaging sessions (Horien et al. 2019), participant cohorts (Dadi et al. 2019), and parcellations (Arslan 2018), while also expressing individual differences (Marek et al. 2019), state-dependent changes (Betzel et al. 2020), and genetic associations (Demeter et al. 2020). Among its characteristic network features is community structure, the presence of reproducible modules consisting of regions that are internally densely coupled, reflecting their coherent and correlated activity over time. These intrinsic connectivity (Damoiseaux et al. 2006), or resting-state networks (RSNs), have become enshrined in the cognitive neuroscience literature, providing a fundamental taxonomy and topographic reference frame for mapping brain/behavior relations (Ito et al. 2017; Uddin et al. 2019). Canonical sets of RSNs have been proposed (Power et al. 2011; Yeo et al. 2011), and their consistent spatial layout has been shown to reflect patterns of co-activation in task-driven fMRI activation studies (Laird et al. 2011). As internally coherent, co-activated, co-fluctuating systems they may be taken to represent ‘building blocks’ of the brain’s cognitive architecture that supports specialized brain function. RSNs are not, however, sharply delineated. As has been noted in early mapping studies (Fox et al. 2005), and later revealed with data-driven community detection and clustering approaches (Power et al, 2011; Yeo et al. 2011), functional connectivity exhibits communities at multiple spatial scales, arranged in an overlapping nested hierarchy (Doucet et al. 2011; Akiki and Abdallah 2019). Furthermore, most cognitive processes do not occur within single RSNs, and indeed may require breaking modular boundaries and dynamic reconfiguration of neural resources, including the network’s nodes and edges (Petersen and Sporns, 2015; Braun et al. 2015; Alavash et al. 2019).

Functional systems or RSNs manifest in long-time samples of resting brain activity – indeed, their reproducibility across imaging sessions sharply increases with the length of time samples, leveling off at time scales of tens of minutes (Gordon et al 2017). This raises the question whether RSNs manifest only on longer time scales or whether they also ‘exist’ at shorter time scales. Recent studies of time-varying functional connectivity (tvFC; Heitmann and Breakspear, 2018; Lurie et al. 2020; Kucyi et al. 2018) have addressed the issue, approaching fine temporal structure and dynamics of FC through the use of shorter data samples, e.g. sliding windows or instantaneous co-activation patterns that result in temporally ordered sequences of functional networks and network states (Liu and Duyn 2013; Allen et al. 2014; Shakil et al 2016; Preti et al 2017). These studies have provided evidence for significant fluctuations of functional connections and network communities on time scales of tens of seconds to minutes (Liao et al 2017; Grandjean et al. 2017; Liégeois et al. 2019; Vohryzek et al. 2020; Hilger et al. 2020). These fluctuations involve a shifting balance between segregated (high modularity) and integrated (low modularity) states (Zalesky et al. 2014; Betzel et al. 2016), with episodes of high modularity exhibiting consistent topology across time (Fukushima et al. 2018) and subject to modulation by task performance or behavior (Shine et al. 2016; Cohen 2018).

Recently, we suggested a new approach to functional connectivity, by focusing on the dynamics and networks formed by ‘edge time series’ (Faskowitz et al. 2020; Esfahlani et al. 2020). The approach unwraps time-averaged FC into time series of co-fluctuating signals on network edges resolved at the time scale of single frames in MRI acquisition, thus allowing inspection of network dynamics at fine time scales. Here we build on this approach and show that a simple proxy for the resulting framewise community structure, expressed as a set of bipartitions of the network into two positively co-fluctuating ensembles of nodes, represents a compact decomposition of the full functional connectivity. Examining the patterns and frequencies of these bipartitions allows addressing several issues related to FC dynamics. How well do bipartitions, sampled at fine-scale temporal resolution, represent ‘classic’ system-level architecture as derived from long-time FC? How does the community structure of single frames combine into the community structure of FC? Do systems, as coherent blocks, manifest continuously or do they appear transiently at short time scales? Do systems differ in their patterns of ‘functional expression’ across time?

## Results

### Extraction of Bipartitions from Time Series

This expository section introduces the basic constructs employed in this study (Fig. 1); for more detail see Methods.

**Figure 1:**
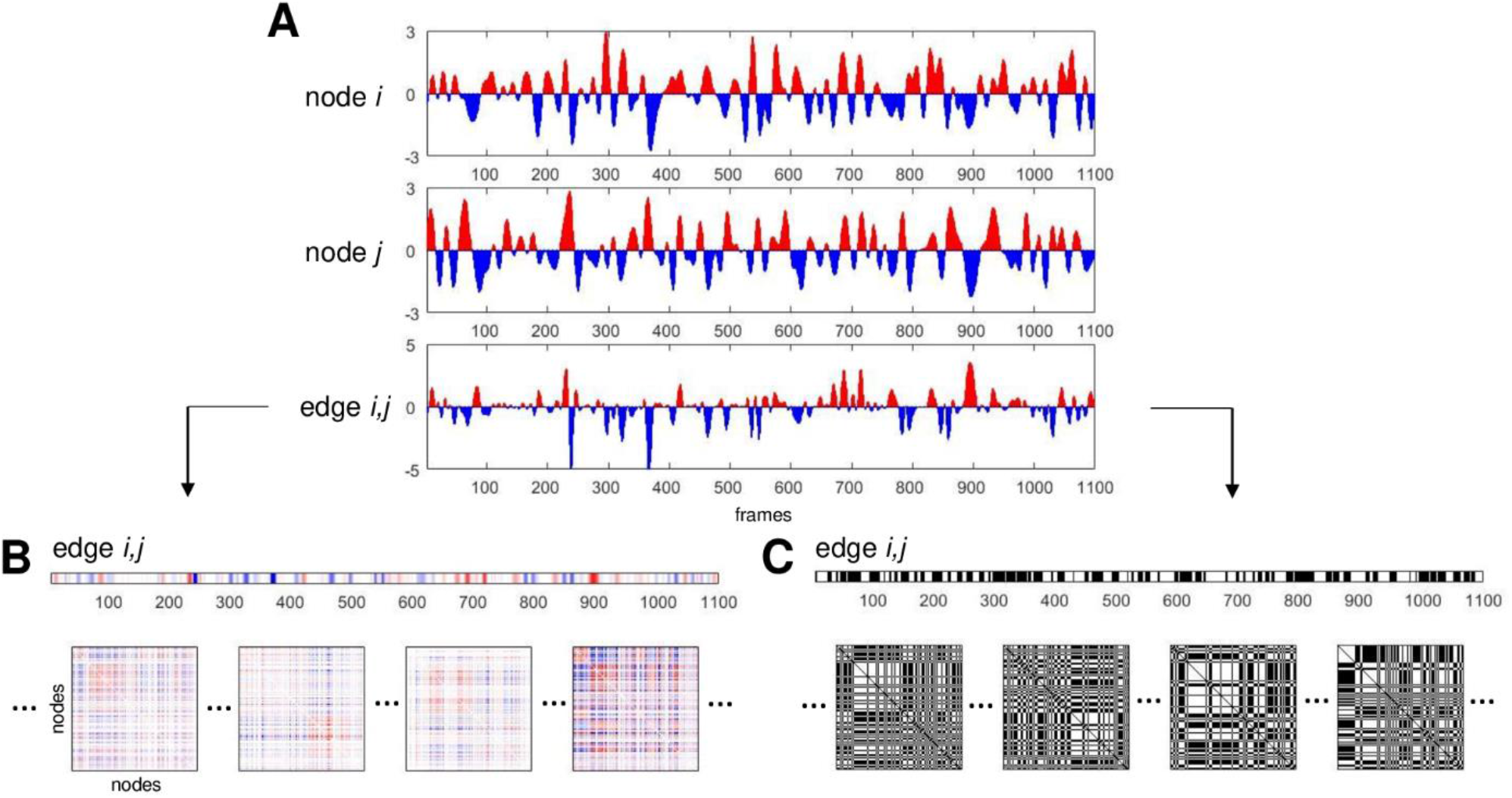
Schematic illustration of main constructs related to time series and bipartitions. (A) Two node time series (BOLD signals, converted to standard scores, for nodes *i* and *j*) and the corresponding edge time series (BOLD signal co-fluctuation, computed as the product of the two node time series, for edge *i, j*). Positive (negative) BOLD signals and positive (negative) co-fluctuations indicated in red (blue). (B) Edge *i, j* time series (same as in panel A) depicted as a matrix row. The set of all (*N*^2^ − *N*)/2 edge time series for a given network composed of *N* nodes can be folded into *N* × *N* matrix form. Examples of single time steps (frames) of such *N* × *N* edge co-fluctuation matrices are shown at the bottom of the panel. The time-average of these single frame matrices is the network’s functional connectivity. (C) Binarized edge *i, j* time series, by thresholding co-fluctuations at *Z* = 0. Positive elements correspond to time points where nodes *i* and *j* exhibit positive co-fluctuations (i.e. the sign of their BOLD signals agree). Frames at the bottom of the panel correspond to the frames shown in panel (B). Each frame is split into exactly two communities. The time-average of these frames is equivalent to the agreement matrix (consensus co-classification) of these communities.

Starting from node time series (BOLD activations), the cross-correlation between each pair of nodes defines their linear statistical dependence (Fig. 1A). The correlations of all node pairs within a given system are that system’s functional connectivity. Employing a standard definition of the cross-correlation, the average of the products of the standard scores of the two variables, yields scalar correlation estimates. Omitting the averaging step retains the summands, corresponding to a temporal un-wrapping of the scalar correlation estimates into vectors (time series) along each edge (node pair). These ‘edge time series’ represent co-fluctuations of node pairs, which are positive when the sign of the two node’s signal amplitude agrees, and negative otherwise. The average of these edge time series is equivalent to the value of the corresponding correlation (functional connectivity) and, when computed across all edges, is equivalent to the FC matrix (Fig 1B). A useful summary metric aggregates the amplitudes of all edge co-fluctuations, computed as the square root of the sum of their squared values (root sum square), here denoted *RSS*. High *RSS* values indicate that node signals strongly agree/disagree at a given point in time.

Removing amplitudes and retaining only the sign of co-fluctuation along edges naturally partitions the network into exactly two sets of nodes (Fig 1C), one set comprising nodes with positive z-scores and a complementary set comprising the remaining nodes with negative z-scores. This is equivalent to thresholding each frame’s node vector at *Z* = 0. The two sets of nodes internally co-fluctuate positively and exhibit negative co-fluctuations between them, thus defining a bipartition of the network.

Reversing the sign of BOLD amplitudes will retain the exact same co-fluctuation pattern and bipartition; we will therefore disregard the signs of z-scored node amplitudes in further analysis. Note also that applying the *Z* = 0 threshold, while inherent to the computation of FC from edge time series, should not imply functional activation of nodes above *Z* = 0 – it merely indicates that regions are active above or below their own mean.

Bipartitions divide the network into exactly two communities, and, over all time frames, these community assignments can be combined into a co-assignment or agreement matrix. In network science, agreement matrices are often used to represent graded assessments of community affiliation (also called co-classification or co-assignment), for example in consensus clustering (Lancichinetti and Fortunato 2012) and multi-resolution community detection (Jeub et al. 2018). In general, the agreement matrix expresses the frequency with which each node pair is grouped into the same community across many partitions. Here, we calculate the agreement of many bipartitions across many time points.

Bipartitions, as special cases of partitions that bisect the network into two communities, are described by a binary node vector of community assignments. The similarity between two such vectors can be measured with several distance metrics such as the Jaccard distance, the cosine similarity, the variation of information, or the mutual information. Here, we adopt mutual information (*MI*) as the principal metric used for assessing similarity between bipartitions. The other metrics are highly correlated with *MI*, and their application gives qualitatively similar results to those reported in this article.

Variations of the bipartition approach are possible. For example, the zero-threshold dividing each frame into two sets of nodes based on the sign of their z-scored time courses may be modified by adopting an arbitrary threshold *θ*. Another approach is to define two thresholds +*θ* and −*θ* that separate highly positive and highly negative activations from activations near the temporal mean, thus yielding tripartions.

### Bipartitions are Strongly Related to Functional Connectivity

All analyses reported in this article have been carried out on four sessions of resting-state fMRI acquired in a cohort of 95 participants, a quality-controlled subset of the ‘100 unrelated’ Human Connectome Project (Glasser et al. 2013) cohort. After pre-processing and nuisance regression each of the four scan sessions was comprised of 1100 frames (TR = 720 ms, total length 792 seconds). BOLD time courses from cerebral cortex were parcellated into 200 nodes according to a standard template (Schaefer et al 2018). Some variations of MRI pre-processing were explored and are discussed below, including a second parcellation scheme into a finer set of 300 nodes and an alternative nuisance regression strategy that retains the global signal (referred to ‘non-GSR data’). For details see ‘Methods’. To allow division of brain regions into a set of functional systems each network node was assigned to one of seven canonical RSNs (Yeo et al. 2011), comprising the visual (VIS), somatomotor (SOM), dorsal attention (DAN), ventral attention (VAN), limbic (LIM), frontoparietal (FP) and default mode (DMN) systems.

Classic FC is equal to the mean over all frames (time points) of the edge time series (Fig 2A). Edge time series can be converted to binary form by applying a threshold based on the sign of the momentary co-fluctuation, an operation that results in a series of bipartitions (Fig 2B). The agreement matrix constructed from these bipartitions is highly correlated with the corresponding FC matrix (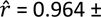 0.008, 95 participants, one session). The strong correlation between this bipartition overlay and traditional FC is robust against different choices of node parcellation and fMRI pre-processing. Fig S1 shows consistently strong similarity between FC and agreement matrix for a finer nodal parcellation (300 nodes) and for time series data omitting global signal regression. Interestingly, for both variants of preprocessing the agreement matrix, after null subtraction, contains negative entries, representing node pairs whose co-assignment into the same module was below chance. In global-signal regressed data these entries strongly overlap with negative functional connectivity. Variants of the bipartition approach also yield high matches of agreement and FC matrices. Adopting an arbitrary (non-zero) threshold *θ* to create bipartitions, or adopting an approach using tripartitions, results in close approximation of FC over a wide range of the *θ* parameter (Fig S2).

**Figure 2:**
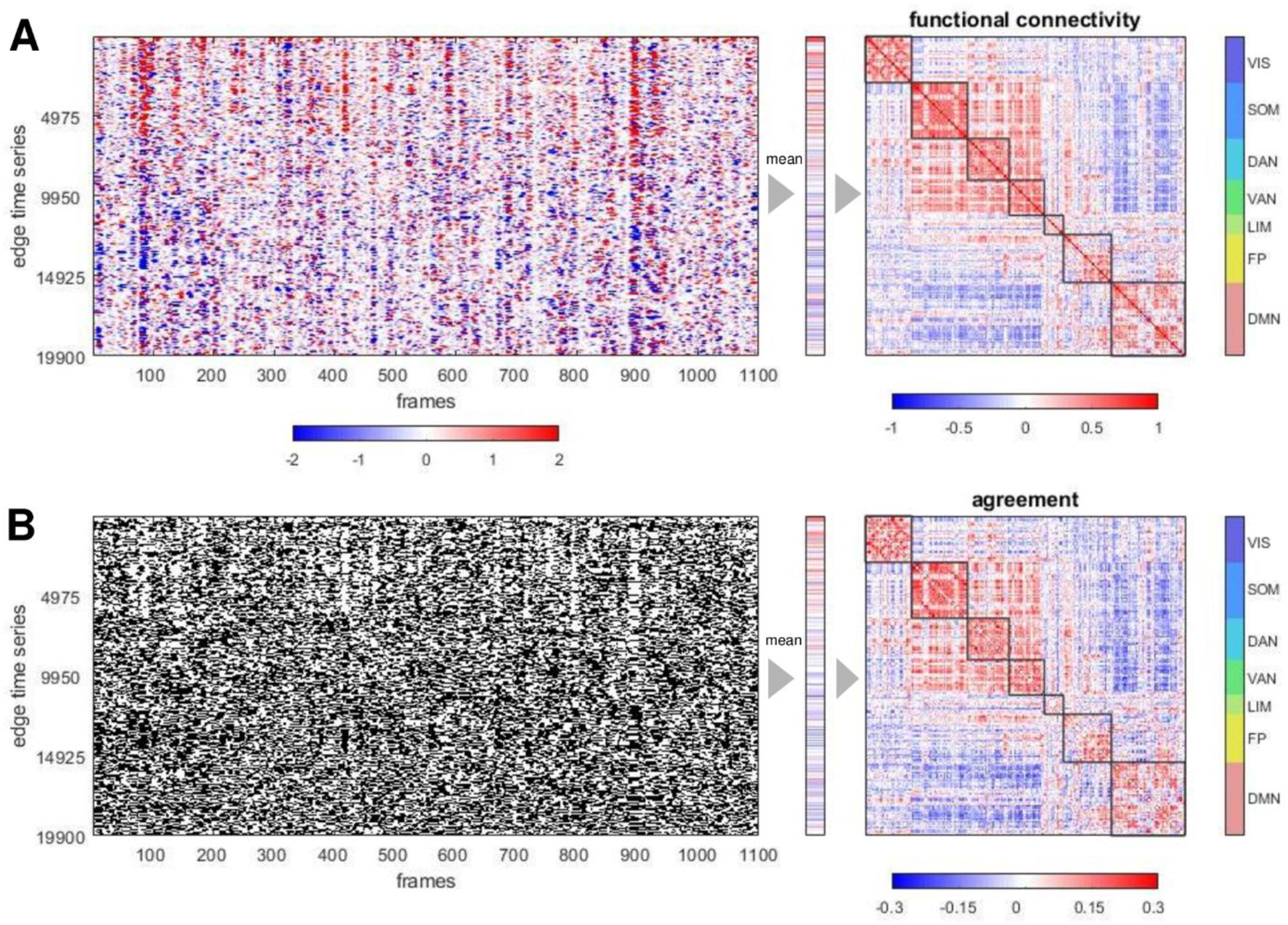
Example of edge time series, FC, bipartitions and agreement matrix, for one representative participant, one imaging session, in a 200 node parcellation of the cerebral cortex. (A) Edge time series recording co-fluctuations between node pairs (19900 unique edges) over 1100 frames (left). The vector of the means of these time series (middle), when refolded into matrix form (right), is equal to the functional connectivity. Nodes are ordered according to 7 canonical functional systems. (B) Thresholding the edge time series at z=0 yields binary time series that track whether co-fluctuations are positive or negative (left). Their average (middle) records, for each edge (node pair), the frequency of positive co-fluctuation which corresponds to their co-assignment (agreement) to the same bipartite community. The agreement matrix (right) is constructed from the complete set of bipartitions and is very highly correlated with the FC matrix (Pearson’s *r* = 0.967, Spearman’s ϱ = 0.963, cosine similarity = 0.962; all computed on the upper-diagonal 19900 element vector).

The set of bipartitions is a binary decomposition of functional connectivity. The characteristic patterning of FC is constructed from the specific spatiotemporal patterns of the constituent bipartitions, as shown in Fig. 3. Subsampling randomly chosen bipartitions gradually approximates FC, with even modest proportions of frames (around 10%) resulting in a very close match with the full-length FC estimate (Fig 3A). When varying run length and using all frames, the quality of the match between agreement and FC matrices remains high even when runs are short (Fig S3). Prior work (Esfahlani et al. 2020) noted that selecting edge time series frames based on their rankings in *RSS* magnitude approximates FC more quickly when frames are ranked from high to low *RSS* amplitudes, as opposed to ranking them from low to high. Bipartitions behave very similarly (Fig 3B). This effect persists when accounting for the autocorrelation structure (temporal adjacency) of the selected frames (Fig 3C). The level to which bipartitions approximate FC is unrelated to framewise head motion. ‘Scrubbing’ (removing) high motion frames (retaining only the frames below the 90^th^ percentile of the framewise displacement) does not significantly affect the match between agreement and FC matrices (Fig S4).

**Figure 3:**
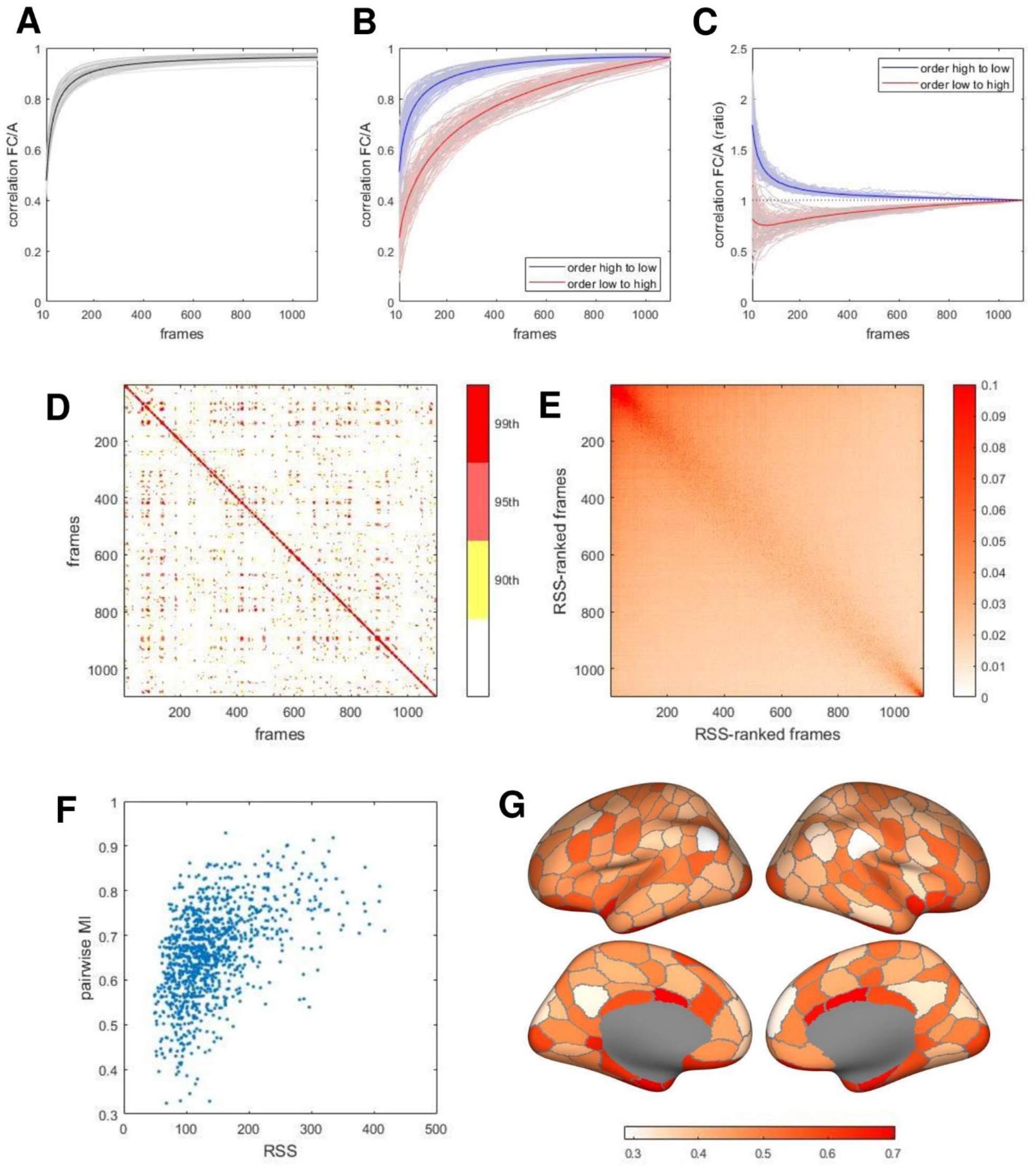
Spatiotemporal patterns of bipartitions. (A) Reconstruction of FC by the agreement matrix constructed from bipartitions as a function of the number of randomly selected time steps (frames). The sample size is varied between 10 and 1100 frames (full length of imaging session), in steps of 10 frames. Correlation of agreement and (full-length) FC is plotted for each of 95 participants (light lines; means shown in black line). (B) Same approach as in panel (A) but with frames selected after ordering them by RMS amplitude. Blue lines show results after selecting between 10 and 1100 frames in descending order of amplitude (going from high to low-amplitude frames), red lines show results after moving in reverse order (going from low to high amplitude frames). (C) Ratio of FC/A correlation when comparing data in panel (B) against a null model (25 independent runs) where frame numbers are shifted by a random offset, thus preserving their number, temporal spacing and hence signal autocorrelations. Ratios greater than 1 indicate better reconstruction than achieved by the null model. (D) Pairwise mutual information between bipartitions on adjacent frames, for a single representative participant and imaging session. Plots displays percentiles of the MI distribution (90^th^, 95^th^ and 99^th^ percentiles). Note recurrent MI peaks between remote frames. (E) Mean pairwise MI, over all participants on a single session, computed after ranking each participant’s frames by *RSS* amplitude. Note high mean MI is predominantly evident on high-amplitude frames. (F) Scatter plot of pairwise MI (adjacent frames) versus RSS amplitude (computed as the mean of the two adjacent frames), in one representative participant on one imaging session. The two measures are significantly correlated (Spearman’s ϱ = 0.502, p = 10^−71^). (G) Switching rates of brain regions, plotted as the ratio of rates observed when *RSS* amplitudes are high versus low. To compute rates, the bipartition communities on selected frames (top or bottom 10% *RSS* amplitude) and their immediate temporal successors were compared to identify those regions that switched their community affiliation. Data were aggregated across all participants and all 4 imaging sessions. The plot shows each region’s number of switches during high *RSS* amplitude frames divided by the number during low *RSS* amplitude frames. All regions’ ratios are less than 1, indicating lower switch rates on high-amplitude frames, with lowest rates exhibited by regions in lateral parietal, medial parietal, and medial frontal cortex (light colors).

Representing the fMRI time series as a series of bipartitions allows computing their pairwise similarity (quantified as mutual information) across time. Fig 3D displays an example of such a similarity matrix for a single participant and a single run. Notably, some instances of bipartitions recur throughout the run as indicated by strongly positive MI between remote time points (off-diagonal entries in the matrix plot). Reordering frames by *RSS* magnitude on each session, followed by averaging over all participants, reveals that high similarity of bipartitions is largely restricted to episodes when *RSS* amplitudes are near maximal (Fig 3E). The bipartition similarity between adjacent time points (pairwise MI) is correlated with *RSS* amplitude (Fig 3F shows data from one representative participant; *ρ* = 0.502, *p* = 10^−71^;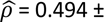 0.049, 95 participants, one session). Lower values of pairwise MI occur when *RSS* amplitudes are small, and higher pairwise MI occurs predominantly when *RSS* amplitudes are large. This relationship indicates that the community structure expressed in frame-wise bipartitions is more stable (changes less) when overall co-fluctuations, across the entire network, are large. These time points correspond to moments when BOLD time series (and hence co-fluctuations), on average, exhibit larger amplitudes, i.e. are farther from their zero mean. Different nodes switch at different rates (Fig 3G), with several DMN regions (parcels ‘DefA_IPL_1’ and ‘DefA_PCC_1’, both hemispheres) and VAN regions (parcel ‘SalVentA_ParOper_1’, right hemisphere) remaining most stably associated with their host communities during high-amplitude epochs.

Principal component analysis (PCA), applied to the set of bipartitions extracted from each participant’s BOLD time course, yields a small number of principal components (PCs) that account for significant portions of the observed variance and exhibit consistent topography across participants (Fig 4A). The largest PC (PC1), on average, accounts for approximately 12% of the variance (12.334 ± 2.082, range 19.989 to 8.780, 95 participants, one session). Averaged across participants and projected onto the cortical surface, the PC1 pattern corresponds to a mode that splits the brain into two co-fluctuating ensembles comprising most regions belonging to the VIS, SOM, DAN and VAN systems on one side versus most regions belonging to the LIM, FP and DMN systems on the other (Fig 4B). The PC1 loadings follow time courses that are positively correlated with *RSS* amplitude (Fig 4C), indicating that the connectivity mode inscribed in PC1 is most strongly expressed at time points with high-amplitude network-wide co-fluctuations. The PC1 as derived from sets of bipartitions is related to several other more familiar constructs (Fig 4D). It is virtually equivalent to the principal eigenvector of FC (or, more precisely, its corresponding covariance matrix), and thus also exhibits strong resemblance to connectivity ‘gradients’ (Margulies et al. 2016). We derived principal components of the affinity matrix computed from FC (equivalent to the principal FC eigenmode) as in the example shown in Fig 4D. The resulting pattern is very highly correlated with the PC1 derived from bipartitions. Furthermore, the bipartition PC1 pattern closely resembles the first principal component of bipartitions identified by applying the Louvain modularity maximization algorithm to the long-time averaged FC matrix (Fig 4D). These strong relationships indicate that the set of bipartitions encapsulates characteristic features of the FC matrix, including its eigenmodes and community structure.

**Figure 4:**
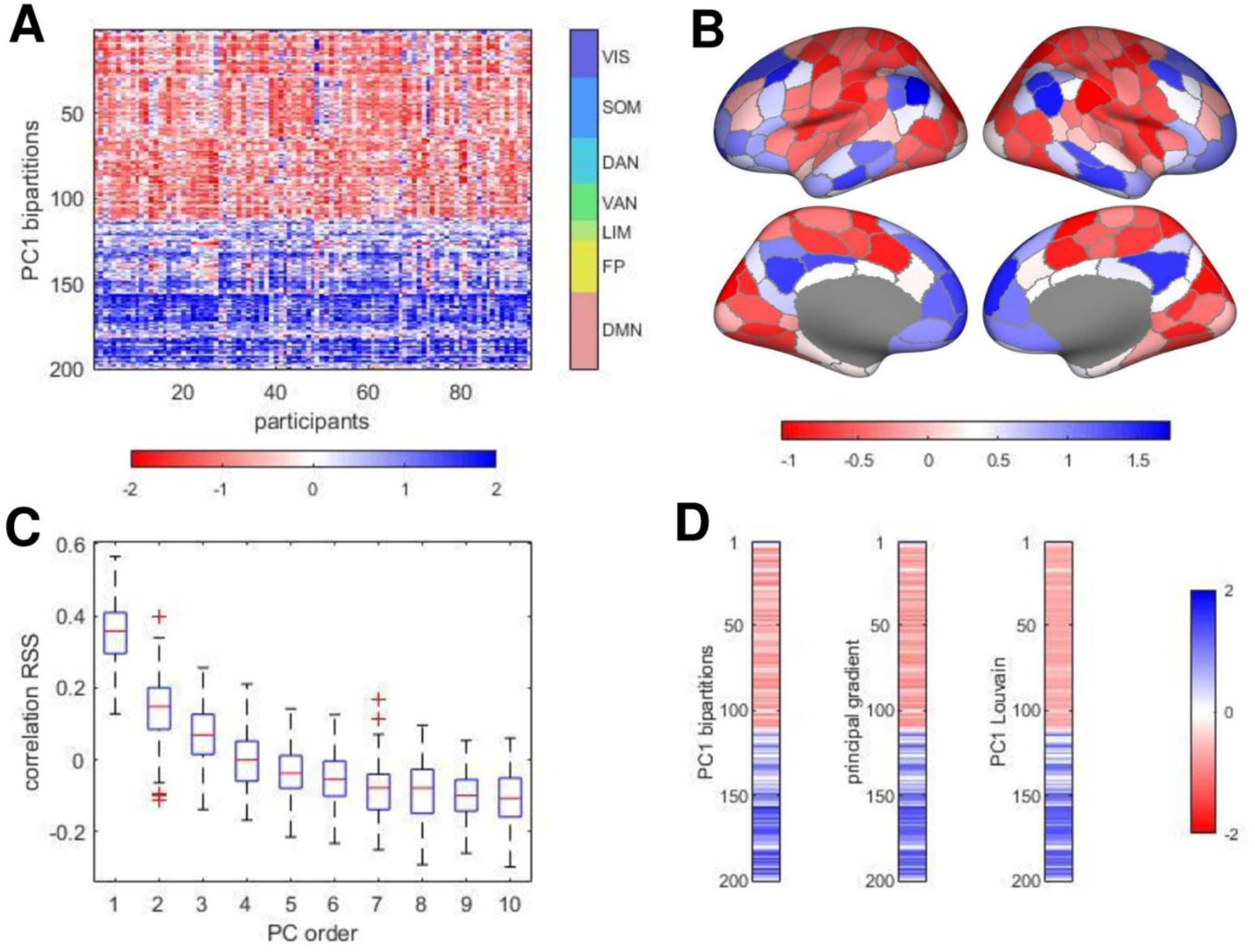
Principal components of bipartitions, and relation to RMS, gradients and Louvain. (A) Largest principal component (PC1) derived from PCA of the complete set of bipartitions, for each participant, single session. All 95 PC1’s are shown, rectified and z-scored to facilitate comparison across participants. The component generally captures a mode that bisects the brain into two sets of functional systems (VIS, SOM, DAN, VAN vs. LIM, FP, DMN). (B) Topography of the PC1 mode (averaged over 95 participants). (C) Boxplot of correlations (Spearman’s ϱ), across participants, of the PC1 loadings, on 1100 frames, with the RMS amplitude computed from the edge time series. Note that components are binned by the order in which they appear in each participant’s PCA but may not directly correspond in terms of spatial topography. Higher-order PCs, accounting for larger percentages of the variance, are more strongly positively correlated with *RSS*. (D) Comparison of the node vectors of the PC1 mode (left), the principal gradient computed from the FC matrix (middle), and the principal component of the PCA of the bipartitions derived by modularity maximization of the FC matrix, using the Louvain algorithm (right). All three vectors represent averages over 95 participants, one session and are z-scored. All pairwise correlations are *r* > 0.98.

Collectively, these results demonstrate a strong relationship between bipartitions and traditional functional connectivity as expressed in the FC matrix. The set of bipartitions derived from the BOLD time series reflects several important spatial and topographic features of FC, while also disclosing its fine temporal structure.

### Bipartitions Map onto Basis Sets of Templates

Bipartitions divide the network, at each point in time, into exactly two communities. These two communities are often approximately equal in size, with only 5% comprising node sets that have fewer than 70 (out of 200) members. This fact begs the question of how these large communities relate to canonical subdivisions of the brain into several, much more compact functional systems, the largest of which (the DMN in the 200 node parcellation) comprising 46 nodes. One way to address this question is to compare each of the empirically observed bipartitions to a standard or basis set of templates that split the brain into bipartitions defined along the boundaries of canonical functional systems (Fig 5A). The basis set used here comprises 7 templates that divide 7 canonical RSNs (Yeo et al. 2011) into 1 versus 6 networks, 21 templates that divide them into 2 versus 5 networks, and 35 templates that divide them into 3 versus 4 networks, for a total of 63 such templates. Since these templates are drawn along RSN boundaries (which themselves are defined based on their coherent co-fluctuations over long time scales) one would expect that bipartitions observed at each frame will at least partially align with the boundaries of the 7 systems as captured in the 63 template basis set.

**Figure 5:**
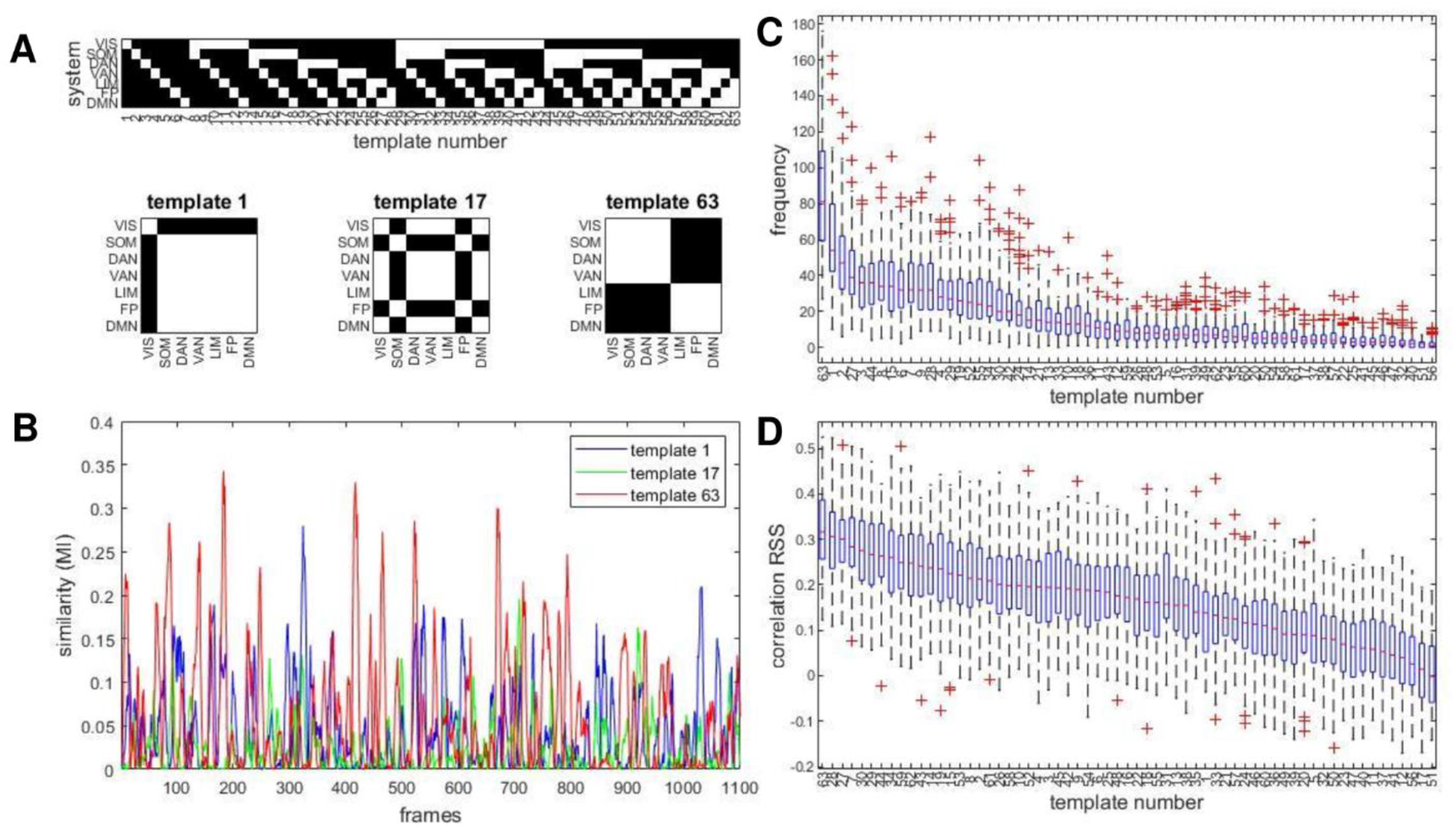
Matching bipartitions to a template basis set. (A) Illustration of the template basis set and examples of templates. Each template is a binary 200×200 node mask, defining a bipartition along the boundaries of 7 canonical resting-state networks. The complete set of 63 templates is indicated at the top. For example, template 1 divides the brain’s 200 nodes into those belonging to the VIS network (29 nodes) and the complement, the remaining 171 nodes. Three example templates are shown at the bottom of the panel. (B) Time courses of the mutual information computed between the observed bipartition and three example templates from the basis set, for a single participant on a single imaging session. (C) Templates that best match observed bipartitions are aggregated across each imaging session and each participant. The boxplot shows their median frequency in order of abundance, across all 95 participants, single session. The frequency is stated as the number of frames when a given template provide the best match (out of 1100 total). (D) Each template’s time course of *MI*, relative to the observed bipartitions on a given session, was correlated to the same session’s *RSS* amplitude. The boxplot shows the median correlation (Spearman’s ϱ) across all 95 participants.

Comparison of templates with observed bipartitions over time allows tracking of several metrics: a) the similarity (mutual information, *MI*) of each bipartition with each basis set template; b) the identification of the single basis set template that most closely resembles the observed bipartition; and c) computing which of these best-matched templates occur most frequently and which correlate most strongly with frame-wise measures such as *RSS* amplitude. Fig 5B shows examples of three *MI* time courses for three examples of templates (cf Fig 5A), one each that divides the network into 1+6, 2+5 and 3+4 systems. The full set of 63 *MI* time courses represent how well each observed bipartition resembles each of the 63 basis set templates and may be interpreted as an index of how strongly a given template is realized at a given point in time. Selecting, at each time frame, the template for which the *MI* is maximal allows representing the sequence of highly variable bipartitions as a sequence of integers, each representing the single best match (highest *MI*) out of the 63 templates. Fig S5 provides examples of observed bipartitions and their best matches in the template set determined by maximal *MI*, for three example templates.

For each participant and scan session, templates can be ordered by their median frequency, based on the number of times they were selected as the best match for the observed bipartitions (Fig 5C). Once a single best-matching basis set template is assigned to each frame, their occurrence can be compared against *RSS* amplitude (Fig 5D). The most frequently observed basis set template (template 63) most strongly correlates with frame-wise *RSS*, indicating that it is predominantly expressed when BOLD signals and their co-fluctuation patterns exhibit high amplitudes. Note that the template 63 pattern strongly resembles the PC1 extracted from observed bipartitions (cf Fig 4A). Qualitatively similar rankings of basis set templates and correlations with *RSS* amplitude are obtained for a finer node parcellation and for non-GSR data (Fig S6).

The best-matching template set represents a highly compressed set of features of the frame-wise decomposition, specific to each imaging session and to each participant. Discarding the temporal ordering of the templates which is immaterial for computing or reconstructing FC, results in a string of 63 numbers encoding a frequency spectrum. Two other aspects of the template set are worth noting. First, the agreement matrix of the template set, as encoded in the 63-element vector, closely matches the down-sampled system-wise FC (Fig S7). Second, the shape of the frequency spectrum across the participant cohort is significantly correlated between imaging sessions. For example, template frequencies for template 63 across all 95 participants is correlated when comparing the mean of sessions 1 and 2 and the mean of sessions 3 and 4 (Fig S8). This correlation suggests that, even after considerable compression of the information contained in the original time series, template frequencies retain some information about individual differences.

### Observed Bipartitions are Constrained by Functional Connectivity

So far, findings indicate that the set of bipartitions observed during single resting-state fMRI runs closely approximates FC (Fig 2) and exhibits characteristic spatiotemporal patterns (Fig 3,4,5). Working backwards from a given FC matrix, we can ask to what extent does the long-term pattern constrain the set of underlying fine-scale bipartitions from which it is composed? Obviously, many different sets of bipartitions (many different sets of time courses) can yield identical FC. To what extent are sets of bipartitions free to vary once their final superposition in FC is fixed?

An optimization approach, searching the space of all possible bipartitions, can help address this question (Fig 6). The approach adopts a variant of the Metropolis algorithm (Metropolis et al. 1953) by maximizing an objective function, defined as the similarity between an empirically observed agreement matrix (which, as established above, very closely resembles FC) and an agreement matrix derived from a set of bipartitions that are subject to incremental optimization. The initial state consists of a completely random set of bipartitions that give rise to a flat agreement matrix. Then, at each subsequent iteration, a single node’s community affiliation on a single time frame (both chosen uniformly and randomly) is swapped. The objective function is re-computed after each swap, and the swap is retained if similarity is increased, subject to a simulated annealing paradigm (Kirkpatrick et al. 1983) applied to ensure that the end state corresponds, as closely as possible, to a global optimum. Three different objective functions are employed, the Pearson correlation, the cosine distance and root-mean-square distance (additional data shown in Fig S9), with near-identical outcomes. Applying the algorithm to data from single participants and single imaging sessions succeeds in identifying sets of bipartitions that closely approximate the agreement matrix derived from the empirical BOLD time series (Fig 6A). Importantly, the optimized set of bipartitions resembles the set of observed bipartitions, as determined by comparing their respective best-matching basis set templates (Fig. 6B). Optimization yields closely matching sets of bipartitions also when the optimized set of bipartitions is significantly smaller than the length of the original time series (Fig S9). For example, if the optimized set is limited to 1/10^th^ of the length of the original time series (110 frames), optimization still converges and resulting bipartitions continue to resemble those in the observed set.

**Figure 6:**
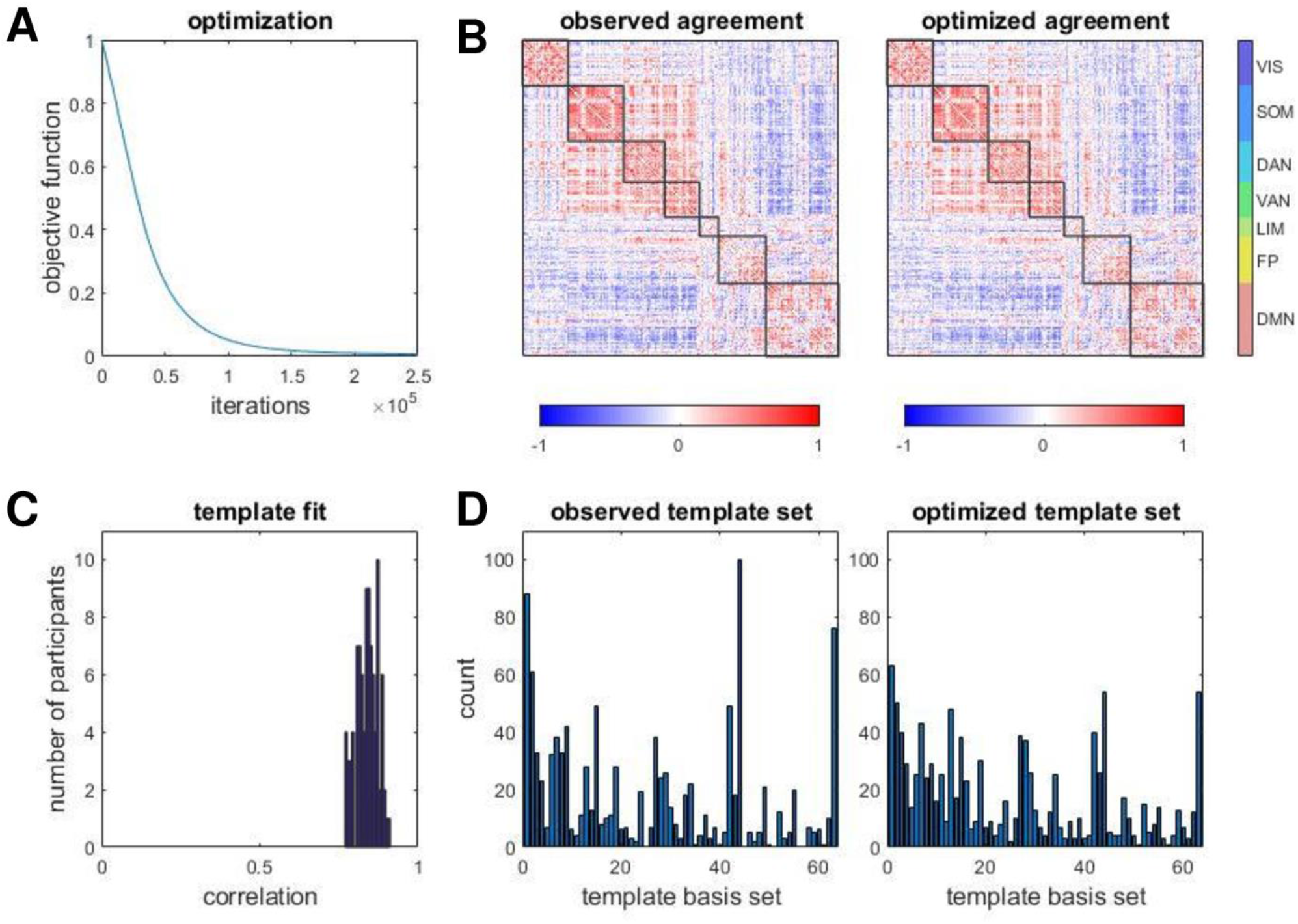
Searching for sets of bipartitions using an optimization approach. (A) Evolution of the objective function (1-Pearson’s correlation) for a single representative participant, single session, single run. (B) Comparison of observed agreement matrix and optimized agreement matrix, the latter retrieved after optimization was terminated (same data as in plot in panel A). (C) Both, observed and optimized sets of bipartitions were compared against the 63 template basis set (cf Fig 4) to retrieve best matches. Their distributions and frequencies were highly correlated (Spearman’s ϱ = 0.840 ± 0.034, range 0.770 to 0.913, 95 participants); correlation magnitudes are shown as a histogram. (D) Example of best matching templates obtained from observed and optimized bipartitions, in a single participant, single session, single run (Spearman’s ϱ = 0.871, p = 0).

These findings suggest that the set of bipartitions encountered in the decomposition of fMRI data is constrained by the long-time average functional connectivity. Recall that each bipartition represents a snapshot of how co-fluctuations distribute across the network, and that the total set of these snapshots exhibits significant fluctuations across time. The optimization approach suggests that these fluctuations are necessary for reconstructing long-time averages in FC, as optimized bipartitions strongly resemble and are as variable as the observed set.

### Expression of Canonical Systems varies across Time

The findings presented so far suggest that bipartitions offer an opportunity to compress time courses into discrete feature sets that retain long-time characteristics of FC while also disclosing fine-scale dynamics. A complementary approach to extract fine-scale network states is possible, as explored in this final section. The expression of individual functional systems across time can be tracked directly, by examining co-fluctuation patterns at fine-scale temporal resolution. The mean co-fluctuation of functional systems can be computed across all 7×7 subblocks (each system and each system interaction), yielding 28 unique time series. An example is shown in Fig. 7A. The temporal averages of these time series are identical to the corresponding down-sampled 7×7 functional connectivity matrix (cf Fig S7). On each time step, mean co-fluctuations are compared to a null distribution derived by randomly shuffling system labels and recomputing co-fluctuations (100 independent shuffles per time step). This comparison yields z-scores for each system and pair-wise system interaction, where the z-score expresses how much the signal deviates from the label-reshuffling null. Discretizing these time courses by applying a z-score threshold yields discrete ‘network states’, with systems and between-system interactions either exceeding or failing to exceed the threshold of expression. Visual inspection of a sample time course (Fig 7B) suggests each of the seven RSNs is significantly expressed, as indicated by exceeding the co-fluctuation z-score threshold, only intermittently, on a fraction of time points. Recall that co-fluctuations should not be taken as ‘mean activation time courses’ as they take on positive values when participating nodes are either jointly above or jointly below their long-time z=0 means.

**Figure 7:**
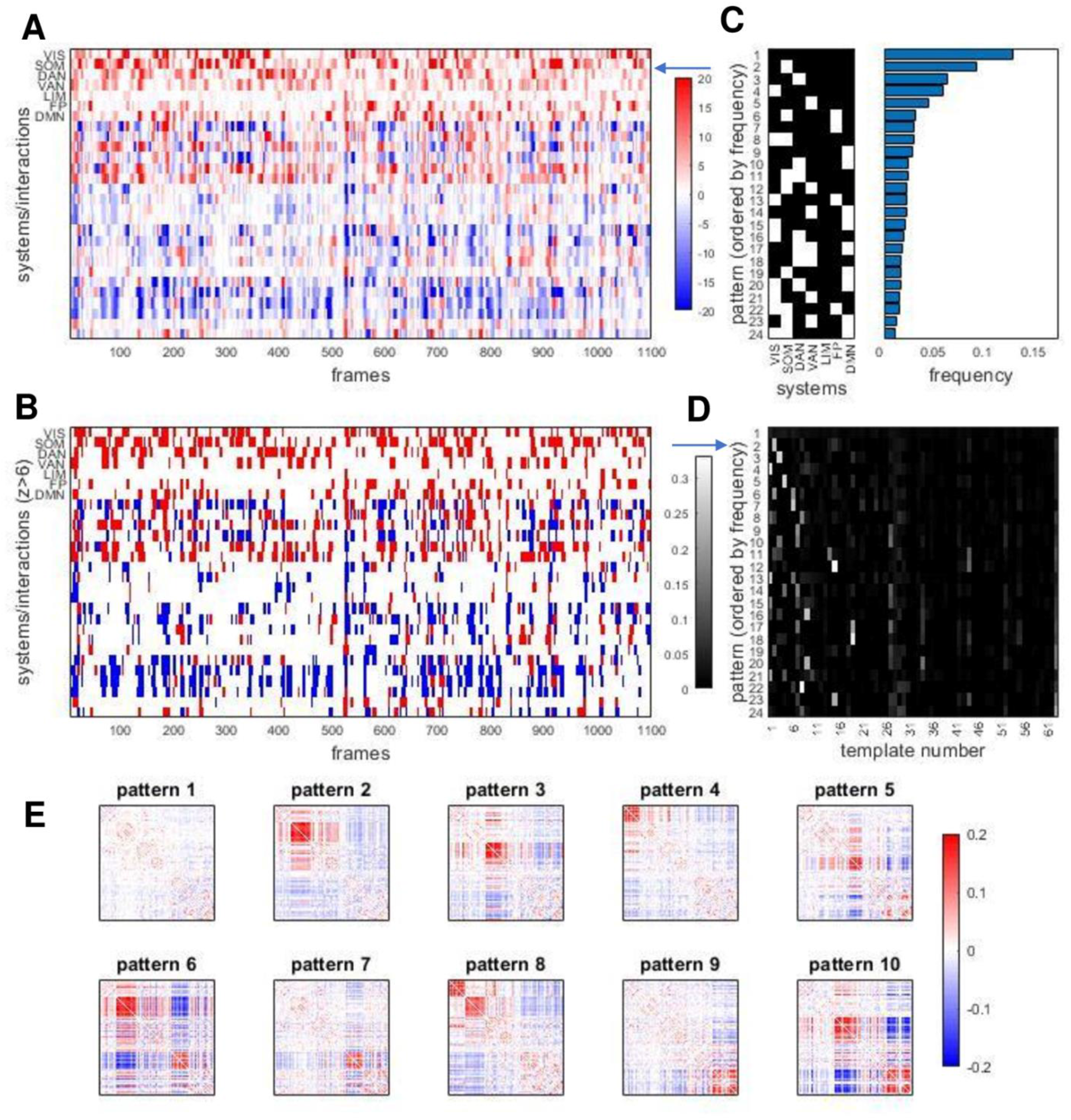
Temporal patterns of RSN expression. (A) Edge time series (cf Fig 2) were aggregated (averaged) based on edges’ placement within or between 7 canonical functional systems. This is equivalent to down-sampling 200×200 (node x node) frames into a 7×7 (system x system) matrix, the latter comprising 28 unique elements. The plot shows the resulting 28 edge time series, for a single participant, single imaging session. Note that within-system time courses exhibit intermittent peaks of high and almost exclusively positive co-fluctuations. Between-system interactions show similar intermittency, with both positive and negative co-fluctuations. (B) Same data as in panel A, after discretizing time courses by applying a threshold after z-scoring against a label permuting null model. The threshold shown here is set at *Z* = 6/−6. (C) Each column (time step) in panel B correspond to a discrete system state. The plot at the left shows the most frequent states encountered after aggregating all 95 participants, all 4 sessions (comprising a total of 418,000 time steps and states). States are displayed by frequency, ordered top to bottom. States with frequencies less than 1 % of total frames are not shown. Frequencies are plotted at the right, in corresponding order. Variants of the plot for different z-thresholds are shown in Fig S10. (D) Relation of system states with best-matching templates form the 63 template basis set. Each row of the matrix is normalized to 1. Note that the most frequent system state (no system strongly co-fluctuating) has no clear correspondence with basis set. Other states correlate strongly with specific basis set templates, establishing a link between bipartitions and system states. (E) Average co-fluctuation patterns computed across frames during which specific system states are encountered (top 10 most frequent states shown).

Considering the above-threshold expression of each of the seven RSNs (leaving aside their mutual interactions, and noting that strongly negative z-scores do not occur) yields, for each point in time, a binary seven-element vector (a total of 128 such states are possible, with between 0 and 7 RSNs expressed at a given time). Aggregating these states (95 participants, 4 sessions, 418,000 frames) provides summary statistics on their frequency (Fig 7C). The most frequent state (occurring in approximately 13% of all frames) is one where no RSN is strongly expressed. Individual participants range between 7.4% and 22.4%, and expression levels are correlated across imaging sessions (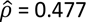, p = 10^−6^, 95 participants, mean of sessions 1 and 2 versus mean of sessions 3 and 4; Fig S10). The next most frequent states predominantly include those where single RSNs are significantly expressed, while states that involve simultaneous co-expression of multiple systems are less frequent. Frequency ranks of states remain stable when changing z-thresholds (Fig S10).

As discussed above, bipartitions decompose FC into sequences of two-community assignment vectors that can be matched to templates from a basis set. As defined in this section, network states also represent sequences of discrete patterns directly derived from significant excursions of edge time series. How do these two representations relate to each other? Network states derived from system-wise expression levels partially reflect the community structure of bipartitions. Many of the states expressing one or several canonical functional systems have clear counterparts within the bipartition template set, i.e. the two representations coincide in time (Fig 7D). Aggregating the bipartitions observed on each time point corresponding to the ten most frequent network states confirms that most states map onto consistent patterns of co-fluctuation as indexed by the bipartition approach (Fig 7E). This comparison establishes a relationship between network states as defined here, through frame-wise averaging of co-fluctuations, and the community structure of bipartitions as defined in previous sections. Both represent compact descriptions of the dynamic expression of functional systems on fine time scales.

## Discussion

Fine-scale analysis of BOLD signal co-fluctuations (edge time series) demonstrates that canonical functional systems are not expressed uniformly or stably across time. Instead their levels of expression fluctuate significantly, as individual functional systems coalesce and dissolve, singly or in varying combinations. While found reliably and reproducibly in long-time scale FC, the appearance of a stable functional systems architecture is the result of the overlap of many transient and fleeting manifestations. The proposed decomposition of FC into bipartitions and networks states allows tracking these dynamics at fine time scales, only limited by the acquisition rate of single MRI frames. The approach complements traditional network analysis of FC, estimated on long time scales) and of time-varying FC, estimated on shorter windows or epochs.

We propose that FC can be decomposed into sets of bipartitions that map onto discrete network states. These bipartitions exhibit characteristic spatiotemporal patterns, with systems and combinations of systems expressed at different times, and in varying combinations. The most common patterns are those where none of the systems are expressed, or where systems are expressed singly and in isolation. The statistics of bipartitions and network states are reproducible across imaging sessions and participants, and do not appear to depend critically on choices made in fMRI preprocessing (e.g. parcellations and global signal regression). Their patterning reflects, and through superposition creates, the complex multiscale community structure of long-time FC, which has been, to this point, the primary target of functional network analysis.

Our work builds on and extends previous investigations of time-varying functional connectivity that has provided evidence for time-dependent fluctuations in functional connections (Chang and Glover 2010) and network patterns and states (Allen et al. 2014; Zalesky et al. 2014; Pedersen et al. 2018; Shine and Poldrack 2018; Lurie et al. 2020). Consistent with prior studies of tvFC our approach reveals spatiotemporal patterns of network-wide co-fluctuations. Notably, we detect a dominant (segregated or modular) connectivity mode that covaries with overall signal amplitudes (*RSS*), appears intermittently over time and exhibits consistent topography (Shine et al. 2016; Betzel et al. 2016; Fukushima et al 2018). Unlike many tvFC studies, our approach does not require defining sliding windows and hence allows tracking system dynamics at higher temporal resolution. The decomposition of the edge time series into discrete sets of bipartitions and/or network states offers not only a highly compressed encoding of system dynamics but also potential new targets for analysis and modeling of both resting and task-evoked fMRI time series data. We note that the decomposition approach presented here is closely related to other methods, including CAPs and iCAPs (Liu and Dyun 2013; Karahanoğlu and Van De Ville 2015; Liu et al 2018), which measures instantaneous patterns of co-activation. Our method is distinct in that it does not require the user to specify a seed region or a threshold for an ‘event’ (Tagliazucchi et al. 2012; Petridou et al. 2013; Cifre et al. 2020). More importantly, the temporal average of edge time series generated by our approach is exactly equal to time-averaged FC, making it possible to measure the precise contributions of individual frames to the static correlation pattern.

The topography of the dominant principal component of the bipartitions bears strong resemblance to the principal mode of BOLD dynamics observed during high *RSS* amplitude ‘events’ (Esfahlani et al. 2020), as well as patterns characterized by strong excursions (Betzel et al. 2016) or high modularity (Fukushima et al 2018) in time-varying functional connectivity. Similar patterns representing a de-coupling of mainly task-positive from task-negative regions have been described and interpreted in previous studies as a major intrinsic/extrinsic dichotomy in functional architecture (Fox et al. 2005; Golland et al. 2008; Doucet et al. 2011; Zhang et al. 2019). The pattern reported here is also very highly correlated with cortical gradients (Margulies et al. 2016), specifically those derived from eigen-decompositions of the functional connectivity matrix. Indeed, this strong resemblance is due to a mathematical relationship between sets of frame-wise bipartitions described here (a compression of the original time series) and the spatial patterns of FC eigenmodes. Going beyond static patterns such as gradients (see also Faghiri et al. 2019), our approach links these connectivity eigenmodes to fluctuating levels of expression of specific functional systems at fine-scale temporal resolution. Their relation to sequences of cognitive processes, e.g. those underlying ongoing thought (Mckeown et al 2020), is an attractive topic for future study.

We employed an optimization approach to explore whether a given FC matrix can be decomposed into sets of bipartitions that differ radically from the ones that are empirically observed. Our findings suggest this is not the case. Given an observed pattern of FC, the set of frame-wise patterns from which it is composed, or into which it can be decomposed, is not free to vary. Instead, the statistics of these patterns appear strongly constrained by the correlation structure inscribed in long-time FC. Optimizations invariably retrieve sets of patterns that resemble those observed empirically, even though no dynamic generative model is employed. This makes it harder to dismiss the observed frame-wise patterns as artifactual or as massively corrupted by noise or uncertainty. Instead it appears that the fluctuating and variable patterns of observed bipartitions are necessary, in the sense that it is difficult if not impossible to construct the observed pattern of FC from a radically different (‘stationary’, temporally smoother, less variable) set of frames.

This work opens new avenues for future research. The decomposition of BOLD time series into a set of bipartitions and/or network states represents a compression or encoding of the system’s dynamics into a much more compact feature set. Such feature sets may provide novel opportunities for mapping individual differences, relations to demographic or behavioral measures, task-rest reconfigurations, or relation to the underlying anatomy. They may also serve as input for machine learning or multivariate statistical mapping, including those probing the relation of brain to behavior. In addition, we note that the proposed scheme may also apply to brain data obtained with other acquisition methods, including more highly resolved recordings of neuronal populations or individual neurons. The decomposition of FC into framewise contributions allows to selectively recombine subsets of frames to get different patterns of FC. It might be possible to select specific subsets of frames/templates to amplify a brain/behavior correlation.

Limitations of the approach should be noted. As is the case with all studies employing functional neuroimaging, the present work inherits most drawbacks of fMRI methodology including its limited temporal and spatial resolution, the indirect link to underlying neural activity, and measurement noise and statistical biases. Following good practices in data preprocessing, the use of multiple data sources and cautious interpretation of findings can at least partially guard against these limitations. It should also be noted that the basic methodological framework transcends the limitations of fMRI as its mathematical and algorithmic core applies to all time-dependent data sources regardless of origin. Future work should aim to reproduce and refine the proposed feature sets, spatiotemporal patterns, and statistics on system expression by leveraging new data sources and participant cohorts. Interventional studies and multi-modal experimentation are needed to identify putative neurobiological mechanisms that underpin or drive temporal fluctuations.

In conclusion, fine-scale temporal fluctuations in the community structure of resting brain activity suggest that the brain’s functional systems express only transiently, intermittently, and infrequently across time. Their robust manifestation over long-time scales results from the superposition of large numbers of spatially distinct and temporally variable patterns. Novel insights and applications may result from the proposed decomposition of brain dynamics into network bipartitions and states.

## Materials and Methods

### Data Set and fMRI Preprocessing

All analyses reported in this article were carried out on data originally collected by The Human Connectome Project (HCP) (van Essen et al 2013), specifically resting-state fMRI data from 100 unrelated adult participants (54% female, mean age = 29.11 +/-3.67, age range = 22-36). The study was approved by the Washington University Institutional Review Board and informed consent was obtained from all subjects. Participants underwent four roughly 15-minute resting-state fMRI scans (here referred to as four imaging sessions) spread out over a two-day span. For a full description of the imaging parameters and image preprocessing see Glasser et al. (2013). Briefly, data were acquired with a gradient-echo EPI sequence (run duration = 14:33 min, TR = 720 ms, TE = 33.1 ms, flip angle = 52, 2 mm isotropic voxel resolution, multiband factor = 8). Participants were instructed to keep their eyes open and fixate on a cross. Images were collected on a 3T Siemens Connectome Skyra with a 32-channel head coil. Participants were considered for data exclusion based on the mean and mean absolute deviation of the relative root-mean square motion across either four resting state MRI scans (file: Movement_RelativeRMS.txt) or one diffusion MRI scan (file: eddy_unwarped_images.eddy_movement_rms), resulting in four summary motion measures. If a subject exceeded 1.5 times the interquartile range (in the adverse direction) of the measurement distribution in 2 or more of these measures, the participant was excluded. These exclusion criteria were established before the current study commenced. Four participants were excluded based on these criteria. One participant was excluded for software error during diffusion MRI processing. Even though diffusion MRI was not part of the present study, this subset was created to include participants with adequate resting-state and diffusion data for future analysis. The remaining subset of 95 participants have the following demographic characteristics: 56% female, mean age = 29.29 +/-3.66, age range = 22-36. Finally, we note here that we defined framewise displacement as the relative root-mean square motion (file: Movement_RelativeRMS.txt), which was computed with the FSL function rmsdiff via the HCP pipelines. We used this information to censor the resting state scans at a frame-by-frame level in a supplementary analysis.

HCP data were minimally preprocessed as described in Glasser et al. (2013). Briefly, data were corrected for gradient distortion, susceptibility distortion, and motion, and then aligned to a corresponding T1-weighted (T1w) image with one spline interpolation step. This volume was further corrected for intensity bias and normalized to a mean of 10000. This volume was then projected onto the 32k fs LR mesh, excluding outliers, and aligned to a common space using a multi-modal surface registration (Robinson et al. 2014).

A functional parcellation designed to optimize both local gradient and global similarity measures of the fMRI signal (Schaefer et al 2017; Schaefer200) was used to define 200 regions (parcels or nodes) of the cerebral cortex. These nodes can be mapped to a set of canonical functional networks [Yeo]; in the current study we adopt a mapping to 7 canonical networks that comprise the visual (VIS), somatomotor (SOM), dorsal attention (DAN), ventral attention (VAN), limbic (LIM), frontoparietal (FP) and default mode (DMN) systems. For HCP data, the Schaefer200 is openly available in 32k fs LR space as a cifti file. A second processing variant used a finer parcellation into 300 nodes (Schaefer300) following the same basic procedure.

We employed two variants in preprocessing to explore the robustness of main findings reported in this article. All analyses were first carried out on data processed with the inclusion of global signal regression. For this strategy, the mean BOLD signal for each cortical node was linearly detrended, band-pass filtered (0.008-0.08 Hz) (Parkes et al. 2018), confound regressed and standardized using Nilearn’s signal.clean, which removes confounds orthogonally to the temporal filters (Lindquist et al. 2019). The confound regression employed (Satterthwaite et al. 2013) included 6 motion estimates, time series of the mean CSF, mean WM, and mean global signal, the derivatives of these nine regressors, and the squares of these 18 terms. Following confound regression and filtering, the first and last 50 frames of the time series were discarded. Furthermore, a spike regressor was added for each fMRI frame exceeding a motion threshold (0.25 mm root mean squared displacement). This confound strategy has been shown to be effective in reducing motion-related artifacts (Parkes et al. 2018). For validation, we also preprocessed the data using aCompCor (Behzadi et al. 2007). These data were linearly detrended, bandpass filtered, and trimmed identically to the previous strategy. This confound regression included five high-variance signals estimated from the CSF and white matter each (10 total), as well as 6 motion estimates, their derivatives, and the squares of these 12 terms. This strategy did not incorporate spike regressors. Following preprocessing and nuisance regression, residual mean BOLD time series at each node was recovered using Connectome Workbench. All data was visually inspected.

### Functional Connectivity and Edge Time Series

Functional connectivity (FC) is generally estimated from fMRI data by computing the Pearson correlation between the BOLD time series recorded from each node pair. Hence, each FC estimate represents a linear similarity between the respective time courses, interpreted as their mutual statistical dependence. It is, by definition, a non-directed non-causal metric that does not distinguish between node pairs that are structurally (anatomically) coupled or un-coupled. All node pairs maintain nonzero FC, and FC estimates may be negative or positive. In a system comprised of N nodes, the system’s FC matrix has dimensions [*N* × *N*], due to symmetry with a total of *K* = (*N*^2^ − *N*)/2 unique entries (all node pairs *i, j* with *i* ≠ *j*).

One definition of the Pearson correlation coefficient states that it is the mean of the product of the standard scores of the two individual variables. Specifically,

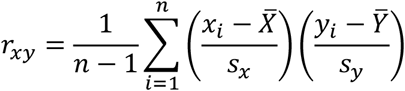

where 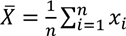 is the mean of x (and applied analogously for y) and 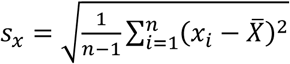 is the standard deviation of x (and applied analogously for y), and *n* is the number of observations (for time series data the number of observations is equal to the number of time points). Thus, there are three steps involved in this computation, the conversion of two node time series to z-scores (note that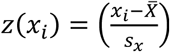), forming their product to create a single time series for node pair *i, j*, and finally forming the mean of this pairwise time series to yield the FC estimate for the node pair (cf Fig 1). This procedure, when repeated for all pairs of nodes, results in a node-by-node correlation matrix, i.e. an estimate of FC. Following an approach developed in prior work (Faskowitz et al 2020; Esfahlani et al 2020; Jo et al 2020), we may omit the final averaging step and retain the time series for each node pair.

Since each node pair subtends a unique network edge, we refer to this construct as ‘edge time series’. The mean of each edge time series is equal to the corresponding node pair’s FC. Rather than collapsing all time steps into a single scalar FC estimate, omitting the averaging step effectively un-wraps FC into a set of edge time series that track ‘co-fluctuation’ between each node pair. At each point in time, the edge time series report a product of two z-scored BOLD signals, which is positive if both signals are above or below their respective zero mean, and negative otherwise. The amplitude of their product varies with the joint amplitudes of the two signals. The full set of edge time series comprises a matrix of dimension [*K* × *T*], with *K* equal to the number of unique edges and *T* equal to the number of time points.

At each moment in time, the amplitude of the co-fluctuations along all edges can be computed as the ‘root sum square’, denoted *RSS*. This metric takes on high amplitude when edge-wise co-fluctuations, on average, are high (either positively or negatively), and it takes on low amplitudes when co-fluctuations are low (again, irrespective of their sign). Prior work has utilized *RSS* to track co-fluctuation amplitudes and stratify or order time points according to their magnitudes. High-amplitude time points coincide with intermittent and recurrent patterns of network activity and connectivity (Esfahlani et al 2020).

### Bipartitions and Agreement Matrix

Edge time series can be converted to binary form, by applying a threshold at the zero crossings that retains only if co-fluctuations are positive or negative. Positive co-fluctuations occur if and only if two signals both exhibit above mean (positive z-score) amplitudes or if both exhibit below mean (negative z-score) amplitudes. Negative co-fluctuations occur when the two signals deviate in opposite directions. A simple extension of this fact is that, on each time step, positively co-fluctuating node pairs split the network into exactly two communities that are fluctuating negatively with respect to each other. This obligatory two-community split results in a bipartition of the network. The network’s time evolution may be represented as a sequence or set of such bipartitions. Adopting bipartitions largely removes information on signal amplitudes (recall that the bipartition does depend on standardizing the individual node time series) while creating a compact description of the original FC as a set of finely resolved modular partitions, without the need to perform computational community detection. Communities are directly evident from the binary edge time series.

It is common practice in network science to combine multiple partitions, for example those obtained from multiple runs of a community detection algorithm, into a single co-classification or agreement matrix (Lancichinetti and Fortunato, 2012; Jeub et al. 2018). The elements of this matrix express, for each node pair, the frequency with which the two nodes are assigned to the same network community. To correct for the rate at which this occurs due to chance, one can subtract the expected frequency if community labels are randomly permuted. Under the assumption that for each sampled partition the number and sizes of clusters are fixed but nodes are otherwise assigned randomly to clusters, one obtains a constant null computed as (Jeub et al. 2018)

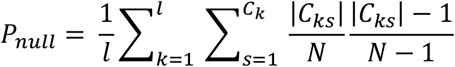

where *l* is the number of samples, *C*_*k*_ is the partition of the *k*-th sample, |*C*_*ks*_| is the number of nodes in cluster *s* of partition *C*_*k*_, and *N* is the number of nodes. In applications to bipartitions from time series, *l* is equal to the number of time points *T*. Subtracting the constant null results in agreement matrices that contain negative entries for all node pairs where the observed frequency of co-classification is smaller than that expected under the adopted null model. The subtraction step does not change relative order of frequencies in the agreement matrix and hence has no impact on correlations or similarity metrics computed against FC.

### Bipartition Similarity

Similarity or distance between two modular partitions can be defined in several ways, including the mutual information computed as

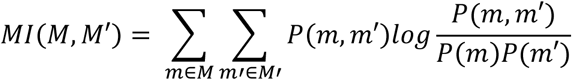

where *M* and *M*′ indicate the two partitions, *m* and *m*′ indicate modules belonging to the two partitions, and 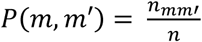 with *n*_*mm*′_ corresponding to the number of nodes that are members of module *m* as well as module *m*′ (Rubinov and Sporns, 2011). In the case of bipartitions, other metrics such as the cosine similarity or the Jaccard distance are also possible. In practice, all these metrics give highly similar results. We adopt the mutual information as the principal metric for assessing similarity between pairs of bipartitions.

### Modularity Maximization

Modularity maximization is a commonly used approach for detecting communities in brain networks (Newman and Girvan 2004; Sporns and Betzel 2016) that attempts to partition a network into non-overlapping communities such that the observed density of connections within subnetworks maximally exceeds what would be expected by chance. The choice of null models should reflect the nature of the network data, which in our case is a correlation matrix (MacMahon and Garlaschelli 2015). We adopt a constant null (the Potts model; Traag et al. 2011) and retain the full FC matrix, including its negative entries, for the purpose of community detection by applying the Louvain algorithm. Louvain bipartitions are identified by first scanning a wide range of the resolution parameter, selecting upper and lower limits within which a two-community structure appears, followed by a finer sampling of this range to retrieve a large set of bipartitions (1000 samples).

### Gradients

So-called gradients, when computed from FC matrices, represent major connectivity modes (node vectors) that can be mapped back onto the original node set, e.g. the surface of the cerebral cortex. Following a standard workflow (de Wael et al. 2020), after starting from an FC matrix, we first derive an affinity matrix that essentially represents a node-wise distance matrix of size [N x N]. Here, we compute the affinity matrix from the full un-thresholded FC as the cosine similarity (1-cosine distance) for each node pair excluding their mutual connections. The first principal component of the affinity matrix is retained for purposes of analysis and comparison. It is virtually identical to the largest eigenvector of the FC matrix. Other variants for computing the affinity matrix may include additional steps such as thresholding or alternative distance transforms. These variants have no impact on the relevant findings reported in this article.

### Optimization

The purpose of the optimization approach pursued in this study was to discover sets of bipartitions of *N* nodes, comprising *P* instances, that approximate the observed bipartitions’ (size [*N* × *T*]) agreement matrix. We adapted a variant of the Metropolis-Hastings algorithm (Metropolis et al. 1953) to generate samples from the very large distribution of possible sets of bipartitions of size [*N* × *P*], starting from a random sample and then iteratively creating variants that are either rejected or accepted, before moving to the next iteration. The rejection or acceptance decision is governed by simulated annealing (Kirkpatrick et al. 1983) and by an objective function *D*, taken here to be a measure of the distance between the optimized and observed agreement matrix. New variants are accepted if the objective function improves (lower distance) or if the annealing criterion *e*^−Δ*D*/*Temp*^ > *R*(0,1) is fulfilled, where *Temp* refers to a simulated ‘temperature’ and *R*(0,1) is a random number uniformly drawn from the [0,1] interval. Essentially, the annealing criterion allows suboptimal variants to pass, as a function of the current temperature. The temperature decays exponentially as a function of the number of iterations, *h*, as *Temp* = *T*_0_*T*_*exp*_^*h*^. Temperature parameters *T*_0_, *T*_*exp*_ were selected such that stable solutions near the global minimum (distance of zero) emerged in reasonable time. The initial conditions were chosen as sets of completely random bipartitions, set with equal probability (‘flipping a coin’) on all node co-assignments. On each step of the optimization, a single element in a single bipartition (both chosen at random) was flipped. Note that this optimization procedure does not implement a true generative process for bipartitions, as they are not derived from time series data and hence contain no information on temporal sequences. While more realistic scenarios for discovering optimally matching sets of bipartitions are conceivable, they were not pursued in the current study.

Three different formulations of the objective function were tested, all computed from the agreement matrix’s *K* unique (upper triangle) elements, with highly reproducible results: (a) the Pearson correlation; (b) the rank-order correlation (Spearman’s ϱ); and (c) the cosine similarity. Optimizations were also carried out by substituting the observed agreement matrix with the observed FC matrix in the objective function, with near-identical outcomes.

The number of bipartitions in the optimized set, *P*, is a free parameter. Different settings of *P*, varying the number of bipartitions from 1100 (matching the number of experimental time steps *T*) down to 11, were explored. Smaller values of *P* yield more compact optimized sets, while also resulting in less accurate matches between the observed and the optimized agreement matrix.

## SUPPLEMENTARY INFORMATION

**Figure S1:**
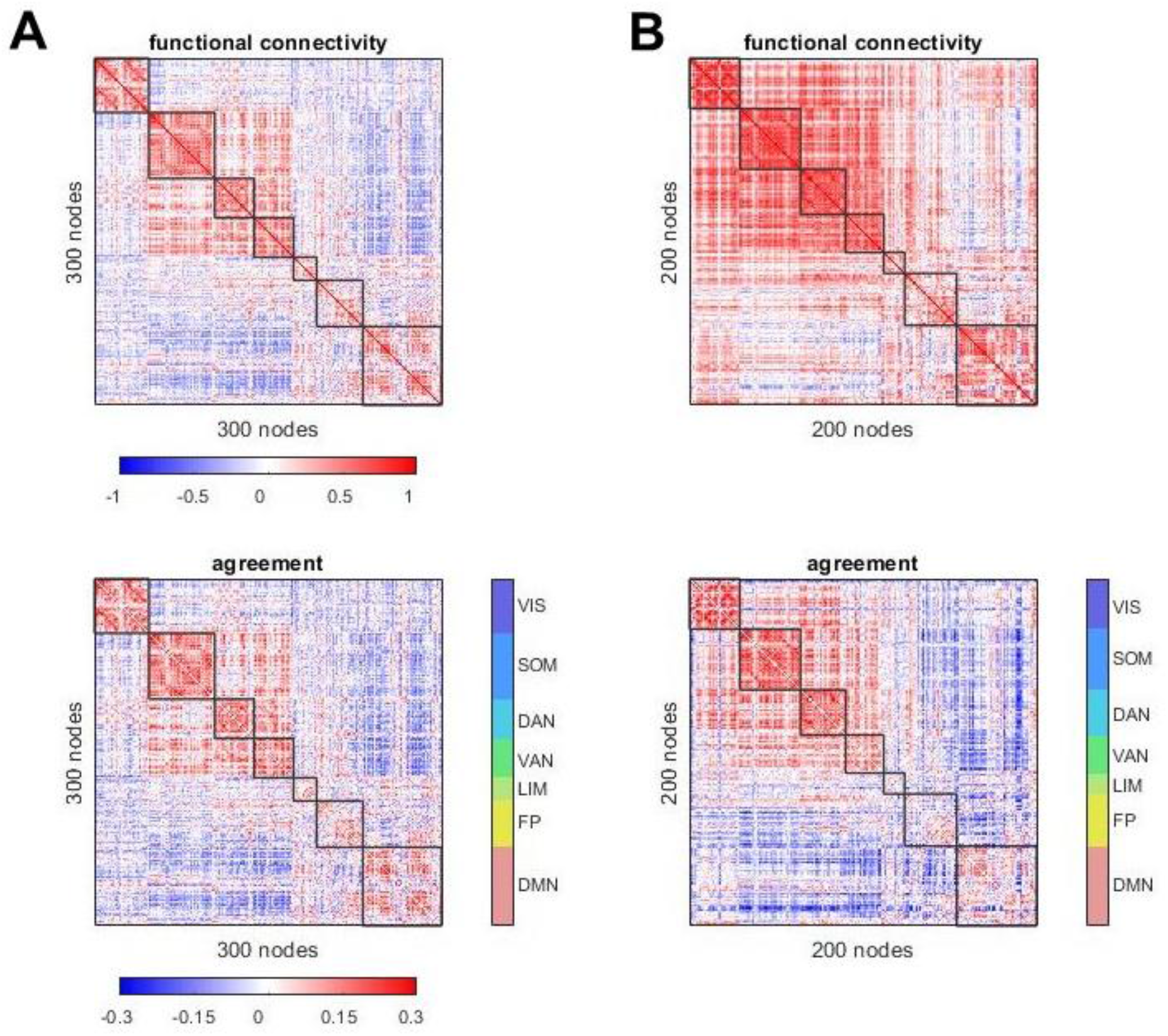
Comparison of FC and agreement matrix, for one representative participant, one imaging session, under finer (300 node) parcellation (A) and when omitting global signal regression from fMRI preprocessing (B). Across all 95 participants the corresponding mean correlations between FC and agreement are ϱ = 0.960 ± 0.008 and ϱ = 0.964 ± 0.009, respectively.

**Figure S2:**
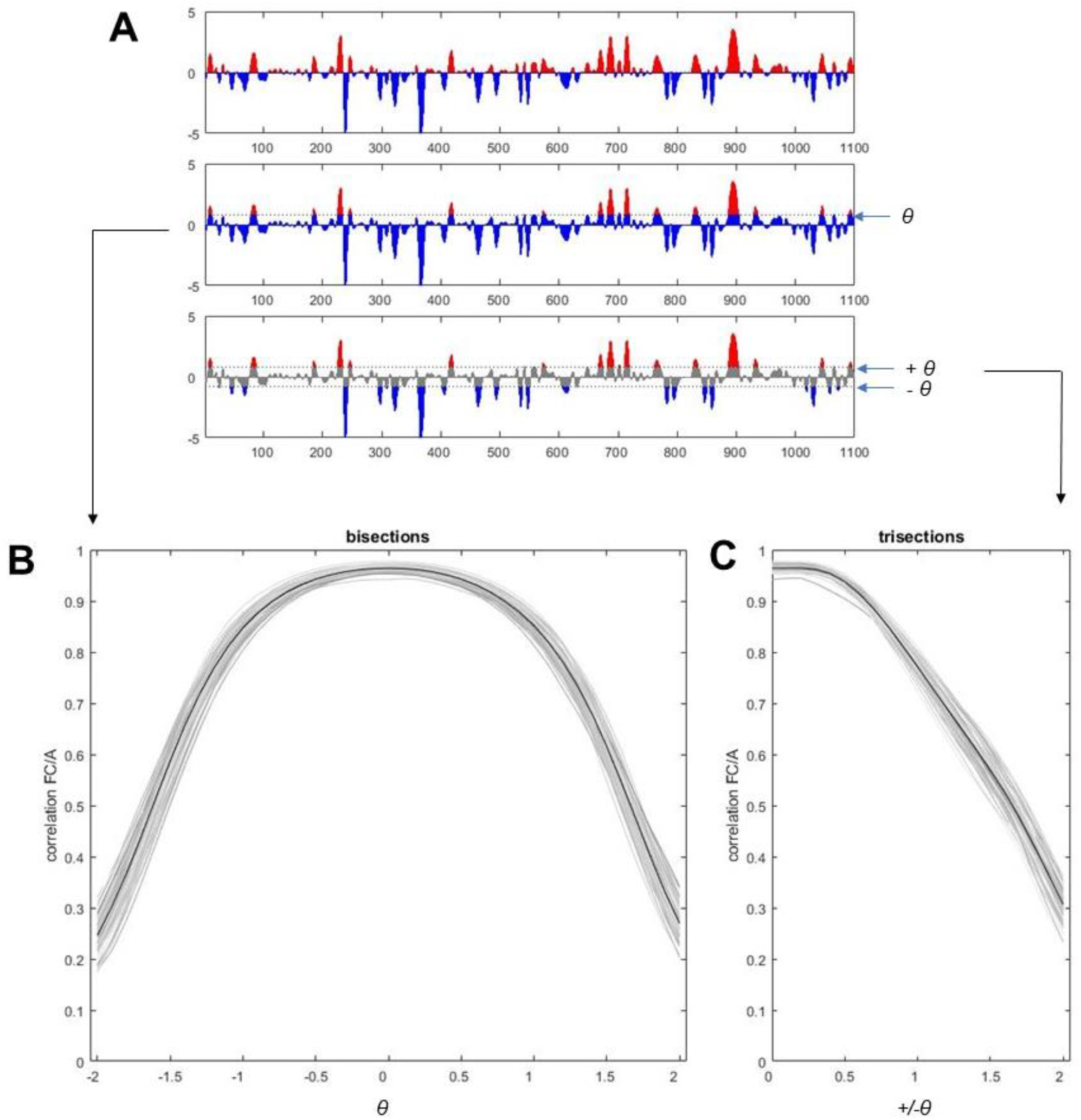
Variant for thresholding to determine bipartitions or tripartitions (A) and impact on correlation between FC and agreement matrices (B,C). (A) Plot at top shows example edge time series (cf Fig 1) thresholded at z=0. Middle panel illustrates choice of an arbitrary threshold ‘tau’ (here tau = 0.8). Bottom panel shows application of two thresholds +tau and -tau to divide the time series into three bins. (B) Correlation (Spearman’s ϱ) as a function of parameter tau, for 95 participants (plot shows individuals as well as group mean). Note that the correlation remains strong over a wide range of the ‘tau’ parameter. (C) Same as panel B, but for tripartitions.

**Figure S3:**
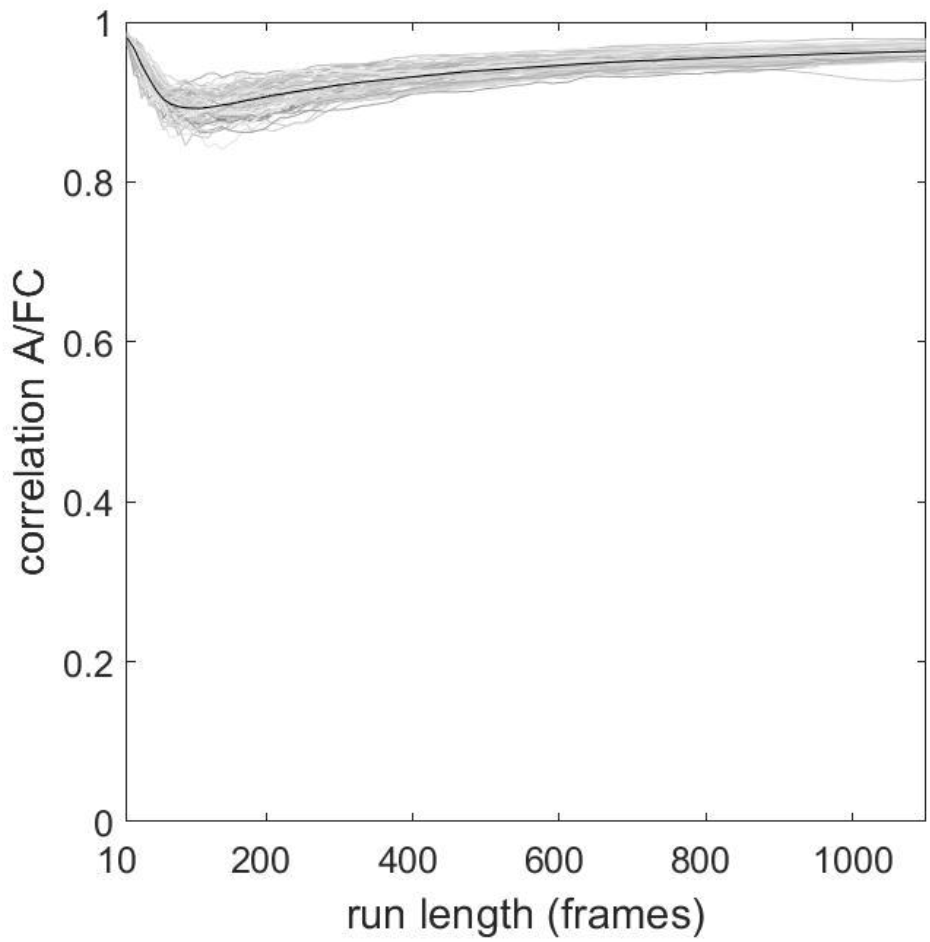
Correlation between FC and agreement matrix as run length is varied from 10 to 1100 frames. FC and agreement matrices were computed for each run length, and thus are both derived from the same length time series data. Even short runs exhibit strong correlations between FC and the corresponding agreement matrix.

**Figure S4:**
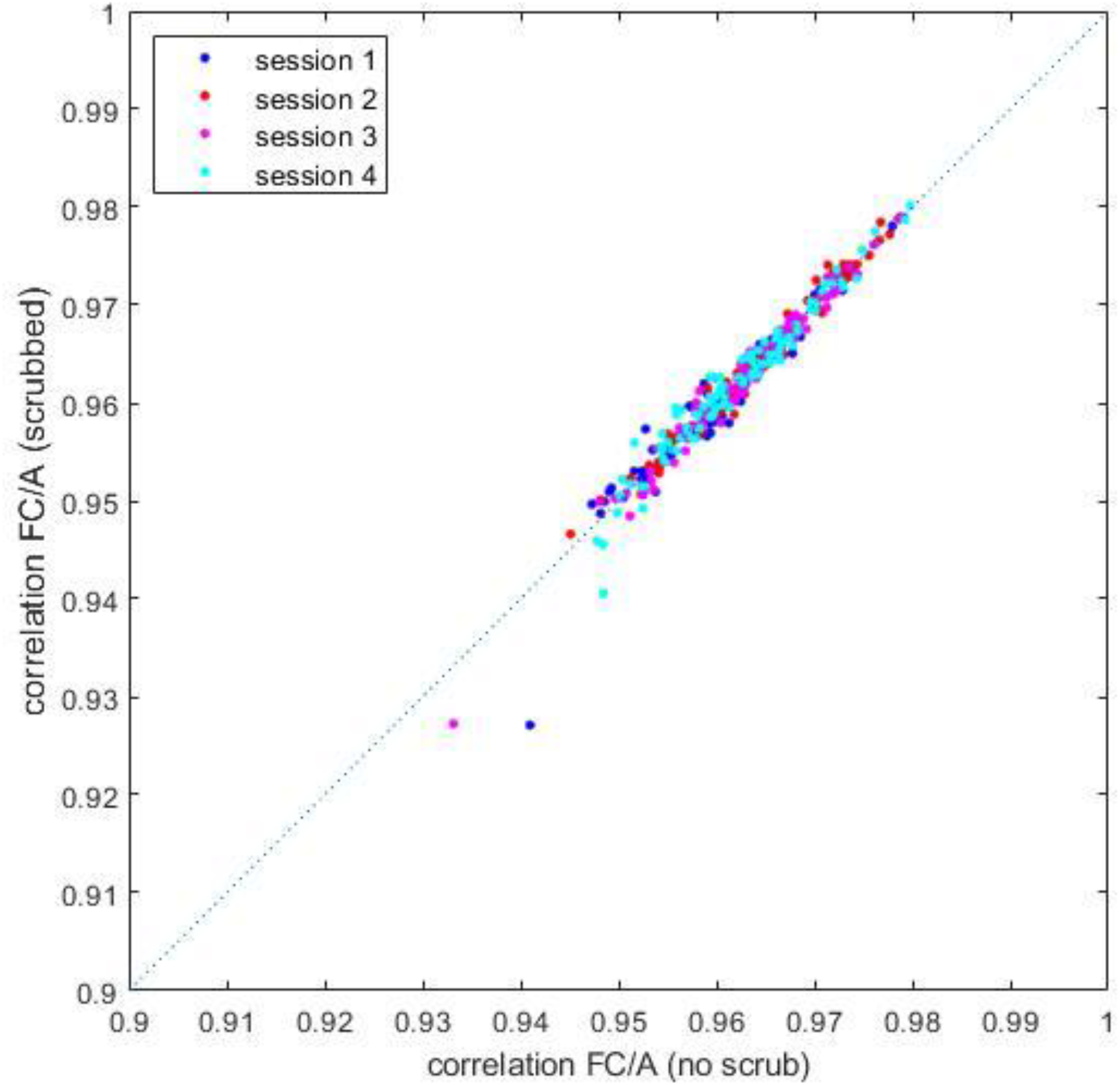
Impact of removing (‘scrubbing’) high-motion frames from each imaging session on the measured correlation between FC and agreement matrix. Scrubbing was carried out by removing all frames for which framewise displacement exceeded the 90^th^ percentile for a given session. The agreement matrix of the remaining 90% of frames was compared (correlated) with the FC (all frames), plotted on the y-axis. The x-axis records correlations of agreement with FC when 90% of frames are sampled at random from the original time series (mean of 100 samples). The plots show data for all 95 participants and for all 4 imaging sessions.

**Figure S5:**
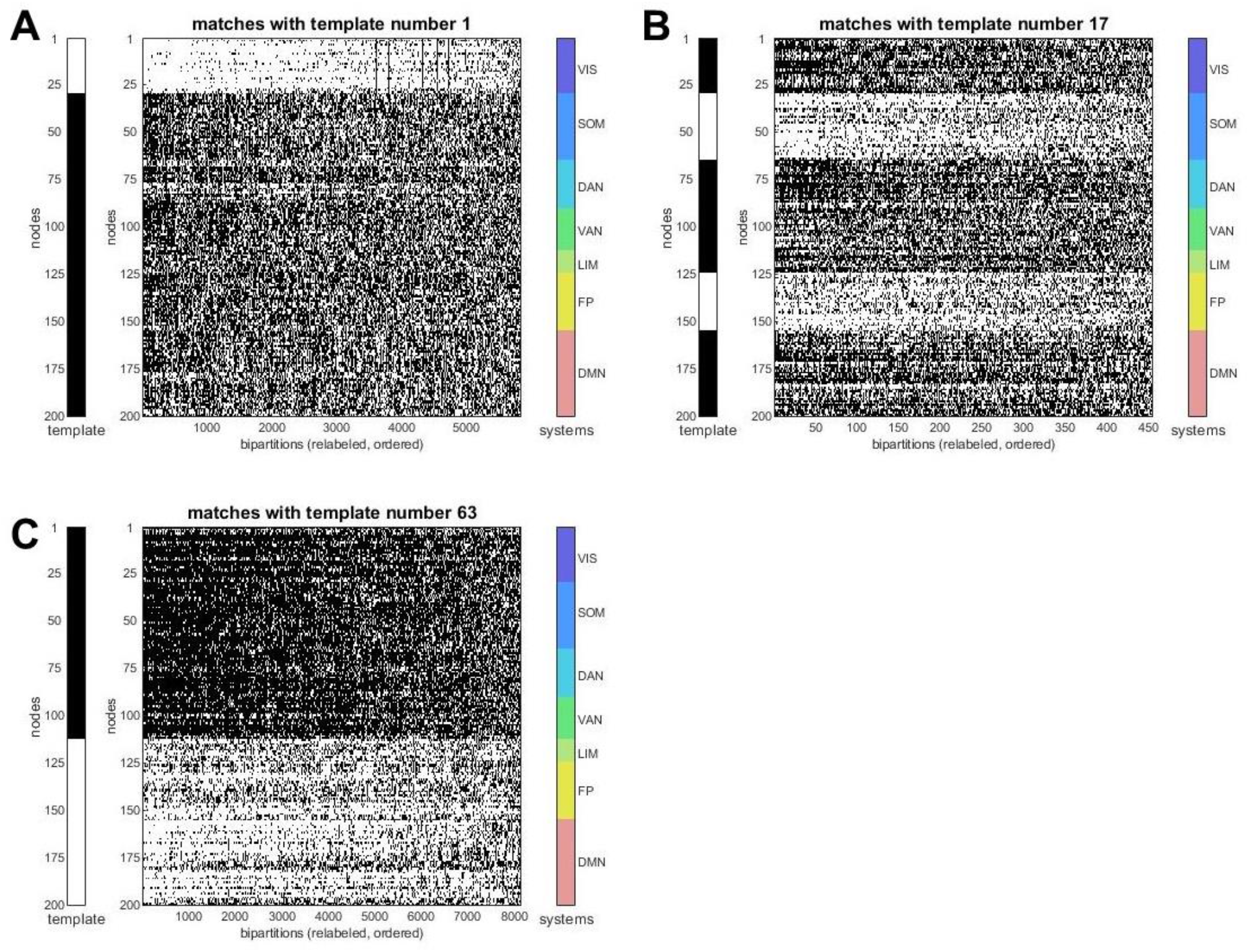
Examples of empirically observed bipartitions that were matched to three different target templates (templates 1, 17, 63; cf Fig 4). Bipartitions have been rectified to facilitate comparison to template vectors (shown to the left in each of the three panels).

**Figure S6:**
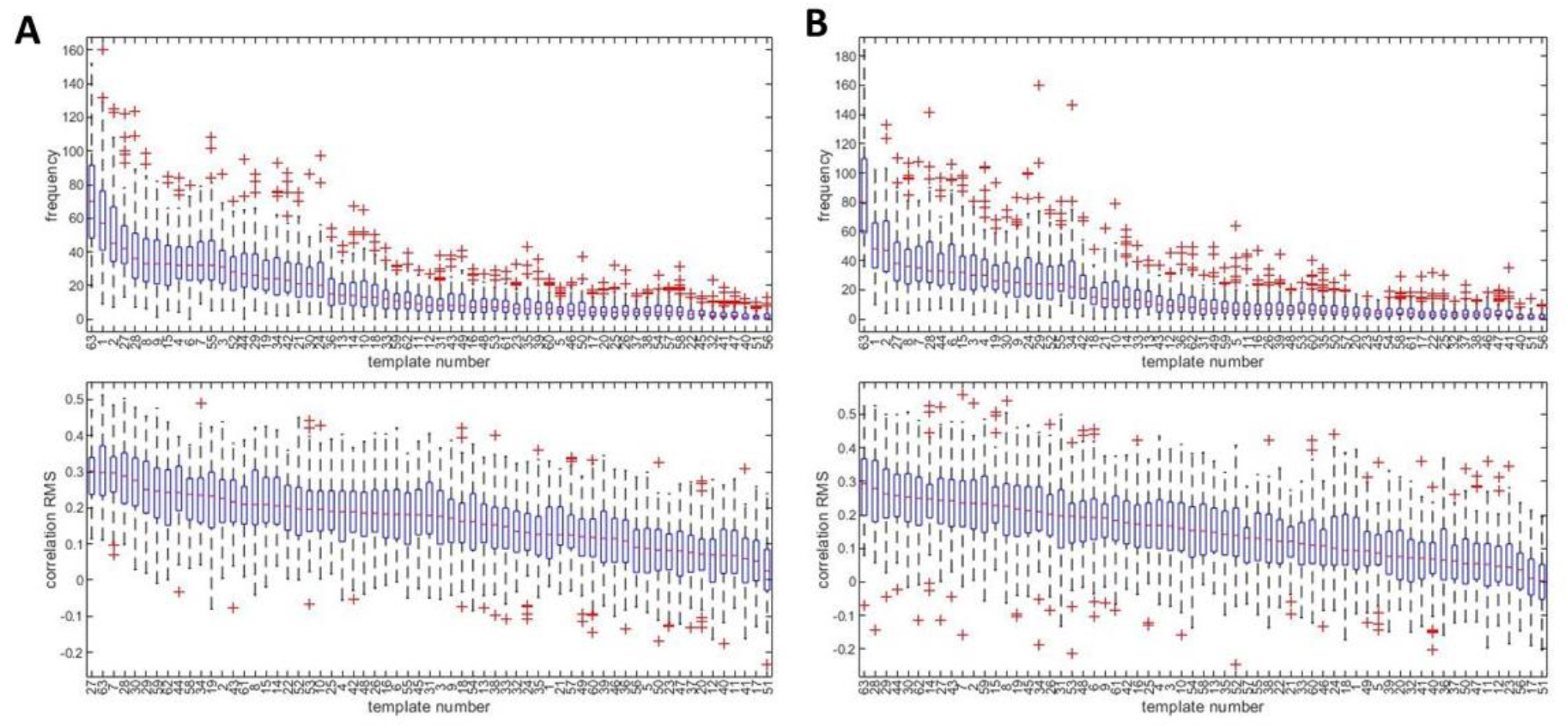
Template frequency and correlation with RSS, for data applying a finer (300 node) parcellation (A) and omitting global signal regression (B).

**Figure S7:**
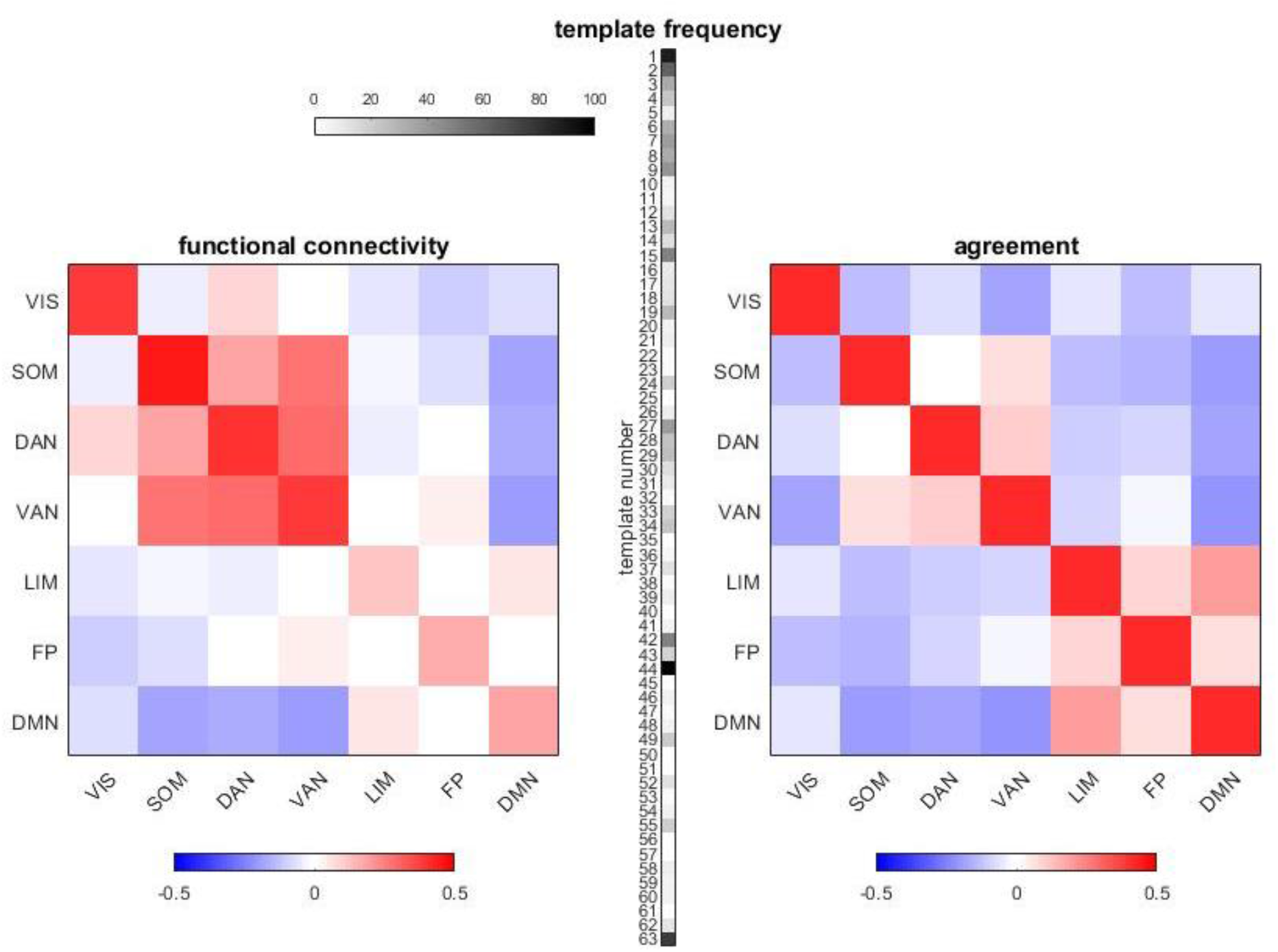
Down-sampled FC (7×7 matrix of functional systems), template frequency vector, and reconstructed agreement matrix, for one representative participant, single session. The template frequency is derived by comparing members of the template basis set to the observed bipartition on each time frame (cf. Fig 4). The agreement matrix on the right is computed from the agreement of the templates as encoded in the vector (middle). Note that structure within systems (matrix elements on the main diagonal) cannot be resolved due to the spatial resolution of the template set. The template frequency vector, which is a highly compressed representation of the original time series, reproduces significant variance in between-system interactions (compare to the down-sampled FC shown at the right).

**Figure S8:**
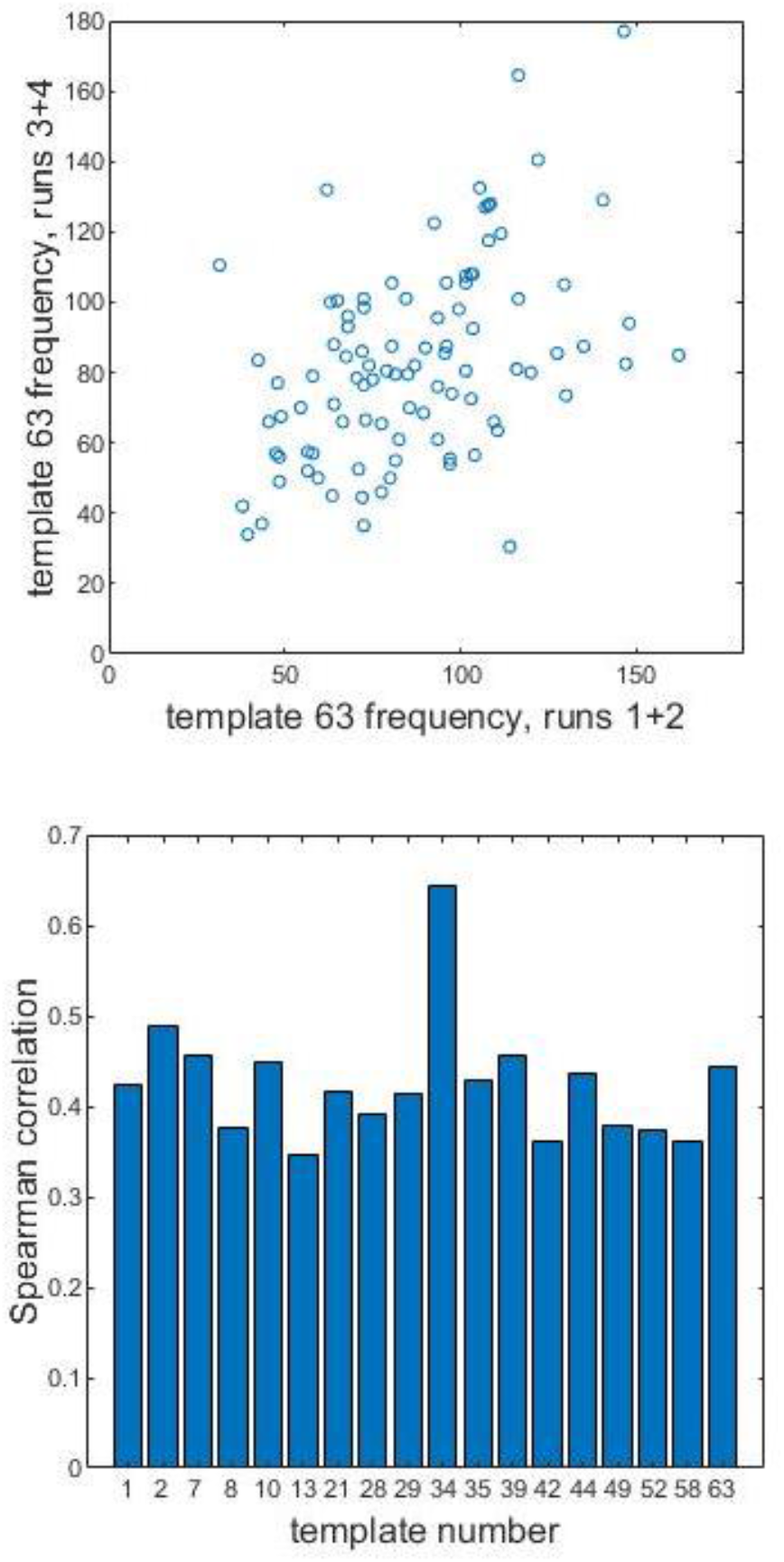
Template frequencies across participants are consistent between imaging sessions. (A) Template frequencies for template 63, per participant, were averaged for sessions 1 and 2 (x-axis) and sessions 3 and 4 (y-axis). Each data point is a single participant. Sessions are correlated with ϱ = 0.444, p = 6×10^−6^. (B) Distribution of cross-session correlations (computed as for panel A) for those templates where the p-value is smaller than the Bonferroni-corrected α of 0.05/63. Note that many of the most frequently occurring templates (cf Fig 4) exhibit significant cross-session correlations. This indicates that some individual differences are preserved in the compressed template representation of individual imaging sessions.

**Figure S9:**
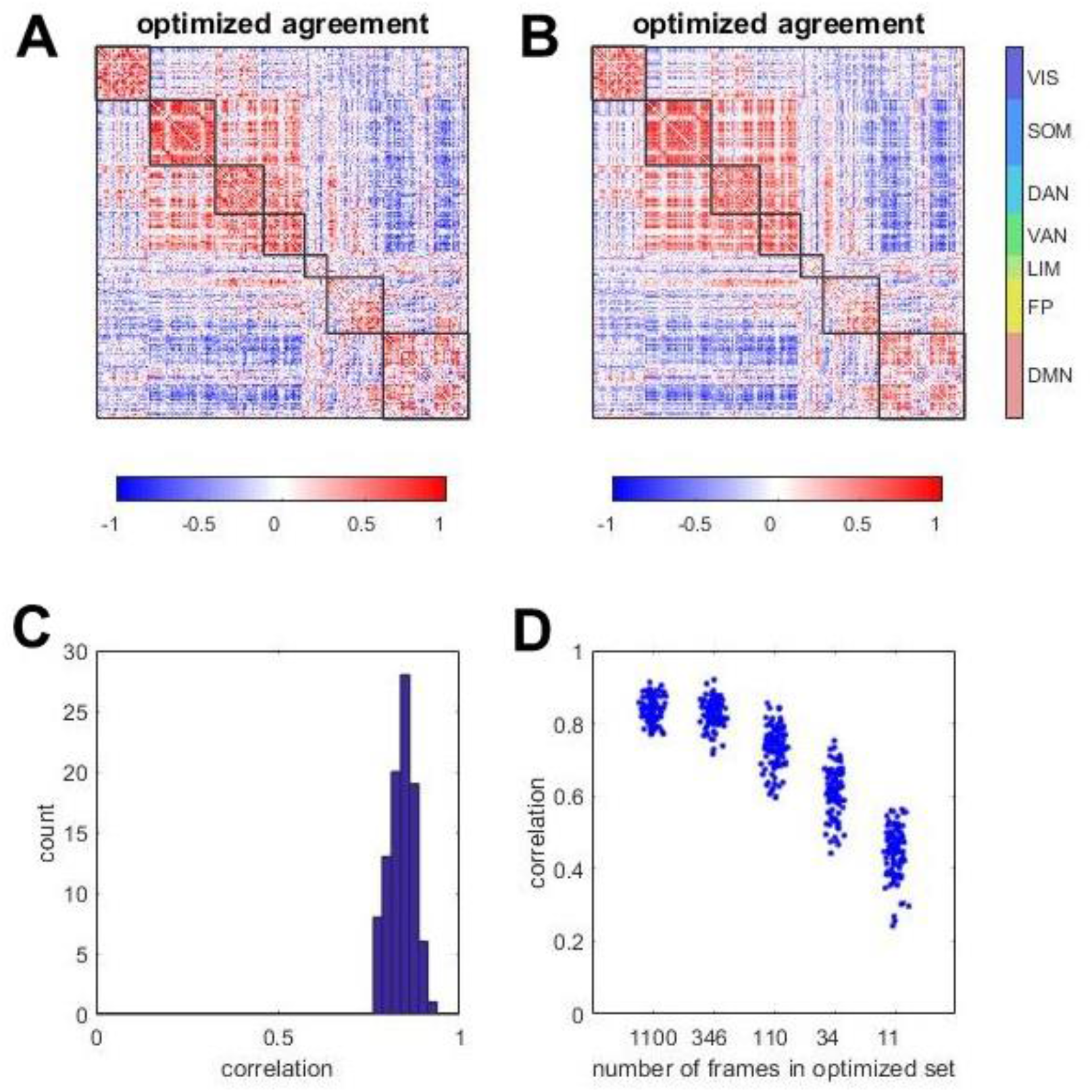
Data from optimizations using Spearman’s ϱ (A) or ‘root-mean-square’ (B) in the objective function. Both panels show optimized agreement matrices for the same participant shown in Fig 6. (C) Histogram of match between template sets in observed and optimized bipartitions, over all 95 participants, single session. (D) Match between template sets in relation to the size (number of bipartitions) of the optimized set. Note that even much more compact sets yield bipartition frequencies that closely match those observed across all 1100 frames in the original time series. In all cases, optimized templates are significantly similar to those found in the observed set.

**Figure S10:**
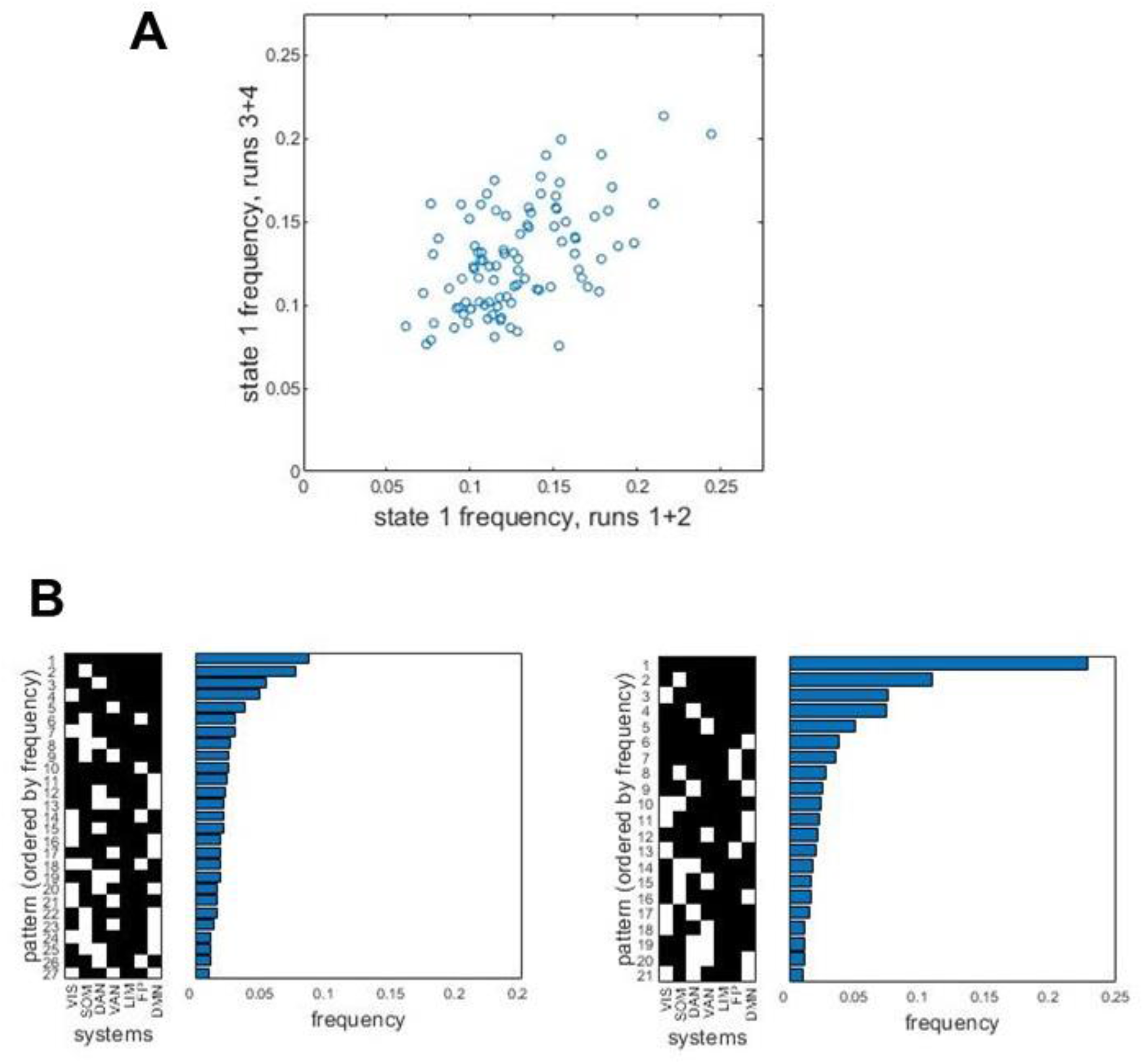
(A) Frequencies of the most abundant system state (pattern 1, cf Fig 7) are consistent across imaging sessions (averages of sessions 1 and 2 on the x-axis, averages of sessions 3 and 4 on the y-axis). Each data point corresponds to one participant. The correlation is ϱ = 0.477, p = 1×10^−6^. (B) Systems states expressed during more than 1% of all frames, ranked by their frequency, when the z-threshold is varied (z = 5, left panel; z = 8, right panel).

